# Intensifying marine heatwaves and limited protection threaten global kelp forests

**DOI:** 10.1101/2024.05.13.594016

**Authors:** Nur Arafeh-Dalmau, Juan Carlos Villaseñor-Derbez, David S. Schoeman, Alejandra Mora-Soto, Tom W. Bell, Claire L. Butler, Maycira Costa, Loyiso V. Dunga, Henry F. Houskeeper, Cristian Lagger, Carolina Pantano, Daniela Laínez del Pozo, Kerry J. Sink, Fiorenza Micheli, Kyle C. Cavanaugh

**Affiliations:** Oceans Department, Hopkins Marine Station, Stanford University, Pacific Grove, California, USA; Department of Geography, University of California Los Angeles, Los Angeles, California, USA; Centre for Biodiversity Conservation, School of the Environment, University of Queensland, St. Lucia, QLD, Australia; Ocean Futures Research Cluster, School of Science, Technology and Engineering, University of the Sunshine Coast, Maroochydore, QLD, Australia; Department of Zoology, Centre for African Conservation Ecology, Nelson Mandela University, Gqeberha, South Africa; Department of Geography, University of Victoria, Victoria, British Columbia, Canada; Department of Applied Ocean Physics and Engineering, Woods Hole Oceanographic Institution, Woods Hole, MA, USA; Institute of Marine and Antarctic Studies, University of Tasmania, Tasmania, Australia; University of Cape Town, Cape Town, South Africa; Universidad Nacional de Córdoba, Facultad de Ciencias Exactas, Físicas y Naturales, Ecología Marina, Córdoba, Argentina; Consejo Nacional de Investigaciones Científicas y Técnicas (CONICET), Instituto de Diversidad y Ecología Animal (IDEA), Córdoba, Argentina; Fundación Por El Mar (PEM), Argentina; Departments of Geography and Anthropology, University College London; South African National Biodiversity Institute, Kirstenbosch, Cape Town, South Africa; Institute for Coastal and Marine Research, Nelson Mandela University, Gqeberha, South Africa; Stanford Center for Ocean Solutions, Stanford University, Pacific Grove, California, USA

## Abstract

Kelp forests are one of the earth’s most productive ecosystems and are at the greatest risk from climate change, yet little is known regarding their future threats and current conservation status. By combining a global remote sensing dataset of floating kelp forests with climate data and projections, we find that exposure to projected marine heatwaves will increase ∼8 times compared to contemporary (2001-2020) exposure for intermediate climate scenarios. While exposure will intensify for all forests, climate refugia emerge for some southern hemisphere kelp forests, which have lower exposure to contemporary and projected marine heatwaves. Under these escalating threats, less than 3% of global kelp forests are currently within highly restrictive marine protected areas, the most effective conservation measure for providing climate resilience. Our findings emphasize the urgent need to increase the global protection of kelp forests and set bolder climate adaptation goals.

## Main

Marine protected areas (MPAs) are a cornerstone of marine conservation^1^. Promoted by international agreements, such as the Convention on Biological Diversity (CBD) Aichi Target 11^2^, the area of marine ecosystems under some form of protection has increased since the turn of the century^3^. Because climate change is the main long-term threat to biodiversity^4-6^, the newly agreed Global Biodiversity Framework at COP15^7^ calls for effectively protecting 30% of the oceans by 2030. A central component of the post-2020 targets is increasing the representation of different habitats under effective protection while adapting to climate change. Although many studies report the protection of critical habitat-forming species, such as corals, seagrass, and mangroves^3^, other essential marine habitats, such as kelp forests, remain largely neglected^8^ (but see^9,10^). Information on kelp forest distribution, threats associated with climate change, and protection status is urgently needed to guide ongoing local and global protection efforts.

Kelp forests dominate >30% of the world’s rocky reefs and are among the most productive ecosystems on earth —comparable to terrestrial rainforests and coral reefs^11-13^. However, marine heatwaves (MHWs) and anthropogenic activities threaten kelp forests^14-17^ and their capacity to provide ecosystem services worth billions of dollars^18-20^. Kelp forests are among the marine ecosystems at greatest risk from MHWs^6^, which is concerning given that MHWs are projected to become more frequent and severe in the next decades^21^. For example, Tasmania and northern California have lost >90% of their kelp forests following MHWs and other impacts of climate change^10,22,23^. Climate adaptation strategies —including MPAs— are urgently needed to halt and reverse this loss^15,24^. Well-managed and highly restrictive MPAs —no-take marine reserves where all fishing activities are prohibited—are the most effective type of MPA for supporting the stability of kelp forests^25^ and their resilience to MHW impacts^26,27^ by facilitating the recovery of higher-trophic-level, which helps control kelp grazer populations and prevent overgrazing of kelp^28-30^.

Monitoring subtidal kelp populations over large spatial and temporal scales can be challenging. However, the largest species (i.e., *Macrocystis pyrifera, Nereocystis leutkeana, Ecklonia maxima*) can be mapped by remote sensing because they create extensive forests that float on the surface. Recent advances in satellite imaging of surface-canopy-forming kelp species provide an opportunity to map their distribution^31^, quantify the threats posed by MHWs, and assess their protection status. These data can also inform other climate-adaptation strategies such as identifying climate refugia^32,33^ —areas less impacted by or more resilient to climate change — for kelp forests. Effectively protecting climate refugia for kelp forests is a priority for conservation^34^ because, in these areas, biodiversity can persist^32^ and enhance the resilience of other kelp forests by maintaining a source of recovery for impacted kelp habitats^24^.

Here, we compile the first comprehensive global map of surface-canopy-forming kelp forests (henceforth “kelp forests”) and leverage these datasets to project the global exposure of kelp forests to MHWs and asses their protection status within MPAs. To create the global kelp forest map, we assemble existing regional and national remote-sensing datasets from Landsat observations (1984-present), supplemented with Sentinel-2 satellite imagery (2015-2019^35^) (Supplementary Table 1; see methods). To project threats to kelp forests from climate change, we estimate future cumulative annual MHW intensities from an ensemble of sea surface temperature (SST) from 11 Earth System models, using three climate scenarios generated under the IPCC Shared Socio-Economic Pathways (SPPs)^36^ (see methods). We then quantify the global protection status and the representation of kelp forests at both country and biogeographic levels (i.e., realm, ecoregions^37^) within MPAs categorized as highly, moderately, or less protected based on restrictions to extractive activities obtained from Protected Seas^38^ (see methods). Our findings reveal increasing threats to all kelp forests from future MHWs, although some southern hemisphere forests may act as climate refuges. We also found that kelp forests remain largely unprotected within restrictive MPAs, the most effective type of MPA, which are poorly represented globally. These findings emphasize the urgent need to increase the global protection and effective representation of kelp forests and, given the scale of the threat posed by future MHWs, for bolder climate adaptation goals for kelp forests.

### Global distribution of kelp forests

We found surface-canopy-forming kelp forests in only 12 nations distributed across 6 biogeographic realms and 32 ecoregions, mostly in mid-latitudes in the Pacific, Atlantic, and Indian Oceans (Fig. 1a). Most of the kelp forests are located in five ecoregions, with 23.7% in Malvinas/Falklands, 20.9% in Channels and Fjords of Southern Chile, 12.8% in Southern California Bight, 10.3% in Kerguelen Islands, and 9.2% in Northern California; while 17 ecoregions combined account for only 1% of the kelp forests (Supplementary Fig. 1).

**Figure 1.**
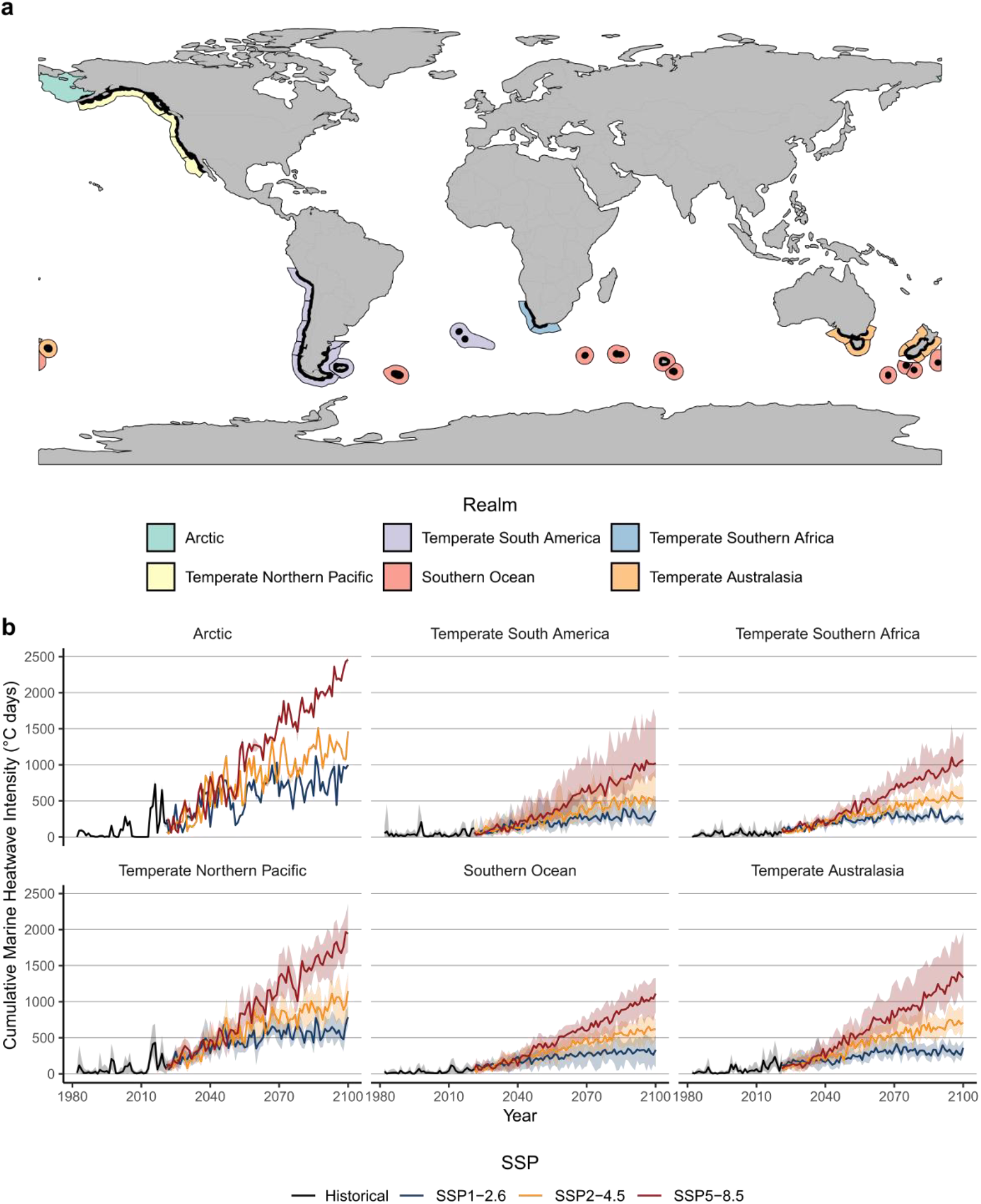
Global distribution of floating kelp forests and exposure to contemporary and future marine heatwaves. Panel **a** map of kelp distribution (black lines) across 32 biogeographic ecoregions (census^37^) (polygons; the color indicates the realm to which they belong), **b** realm-specific exposure of kelp forest to historical (1982-2020) and future cumulative annual MHW intensities (2021-2100) across three climate scenarios (SSP-1.26, SSP-2.45, SSP-5.85). The solid line shows the mean across ensemble medians for all pixels, and the shaded area represents the 5^th^ and 95^th^ percentiles.

In the northern hemisphere, kelp forests can be found at their highest latitudes in the USA (∼61.4 °N), extending southward to their warm-distribution limit in Mexico (∼27 °N). In the southern hemisphere, kelp forests can be found at their lowest latitudes, overall, at their warm-distribution limit in Peru (∼13.6 °S), extending to their highest latitudes in Chile (∼56 °S). Other warm-distribution limits of kelp forests in the southern hemisphere include Argentina, Namibia, South Africa, Australia, and New Zealand.

### Contemporary and Future exposure of kelp forests to marine heatwaves

The exposure of kelp forests to contemporary (2001-2020) average annual cumulative MHW intensities (henceforth “cumulative MHW intensities (°C days)”) was over two-fold higher in the northern hemisphere than in the southern (Supplementary Table 2). The Arctic and Temperate northern Pacific realms were the most exposed to MHWs, particularly the Eastern Bering Sea and the Gulf of Alaska ecoregions, which have registered an average cumulative MHW intensity from 2001–2020 of 177.4 ± 15.1 and 136.7 ± 1.8°C days, respectively. The Temperate South America realm was the least exposed, particularly the Malvinas/Falkland and Prince Edward Islands ecoregions, which have registered an average cumulative MHW intensity from 2001–2020 of only 20.5 ± 0.7 and 23.7 ± 4.3°C days.

Projected future MHWs for kelp forests increase for each realm, ecoregion, climate scenario, and time (Fig. 1b and 2a and Supplementary Fig. 2 and 3). In the near term (2021-2040), kelp forests are projected to be subject to > 2 times higher exposure to cumulative MHW intensities compared to contemporary exposure, with similar values across climate scenarios (Supplementary Table 2-5). Projections suggest that these magnitudes will continue to intensify, and under SSP5-8.5, kelp forests could be subject to > 6 to >16 times higher cumulative MHW intensities in the mid (2041-2060) and long term (2081-2100), respectively, compared to contemporary exposure. These magnitudes are ∼2 to ∼3 times higher than corresponding projections under SSP1-2.6 and SSP2.4-5, respectively.

**Figure 2.**
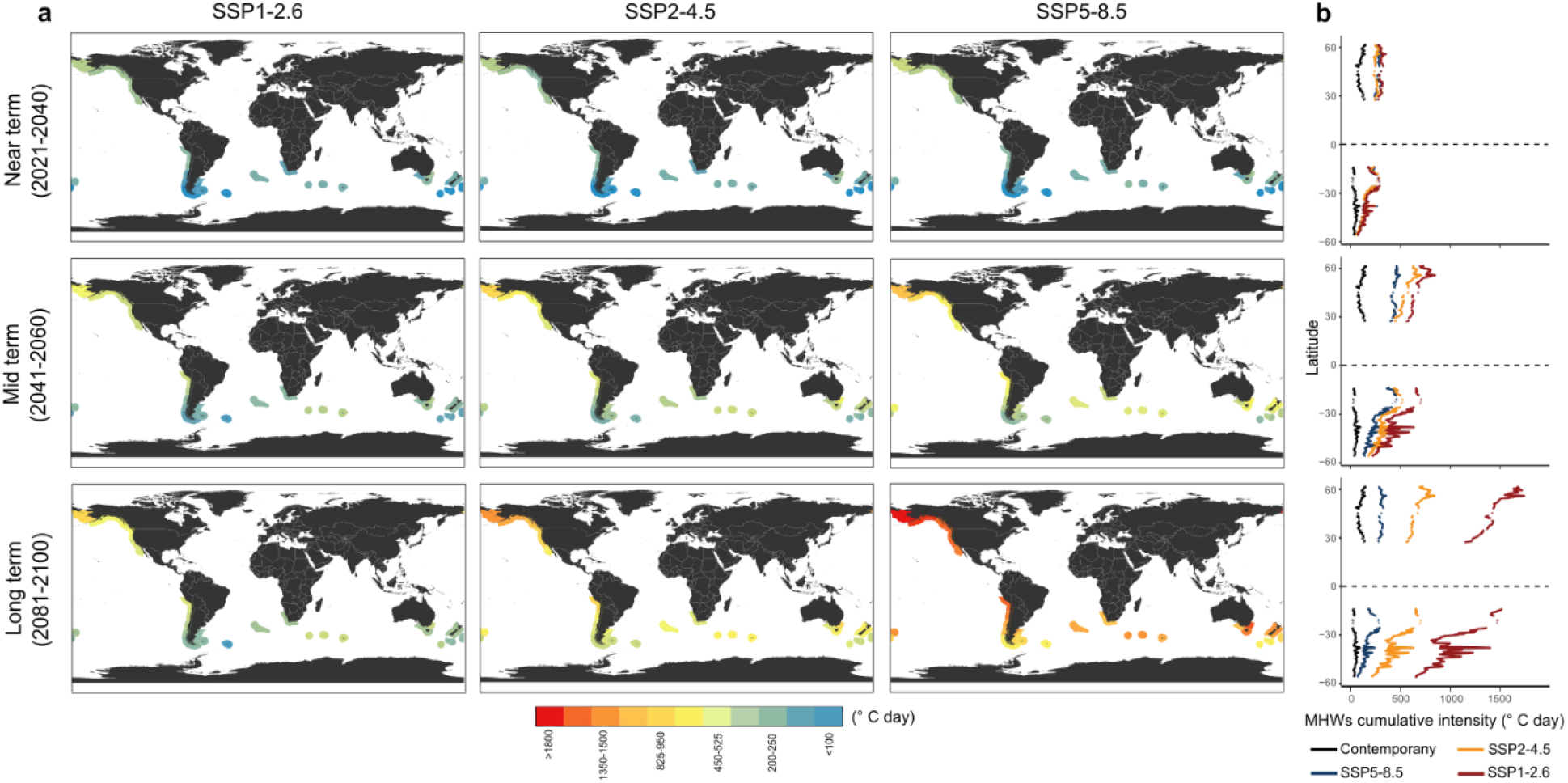
Ecoregional exposure of floating kelp forests to contemporary and future marine heatwaves. **a** mean cumulative annual marine heatwave intensity for all pixels in each of 32 ecoregions under three climate scenarios (SSP-1.26, SSP-2.45, SSP-5.85) and three-time frames (near, mid, and long term). **b** Latitudinal plots representing mean cumulative annual marine heatwave intensities by 1**°** of latitude under contemporary and climate scenarios for each time.

**Figure 3.**
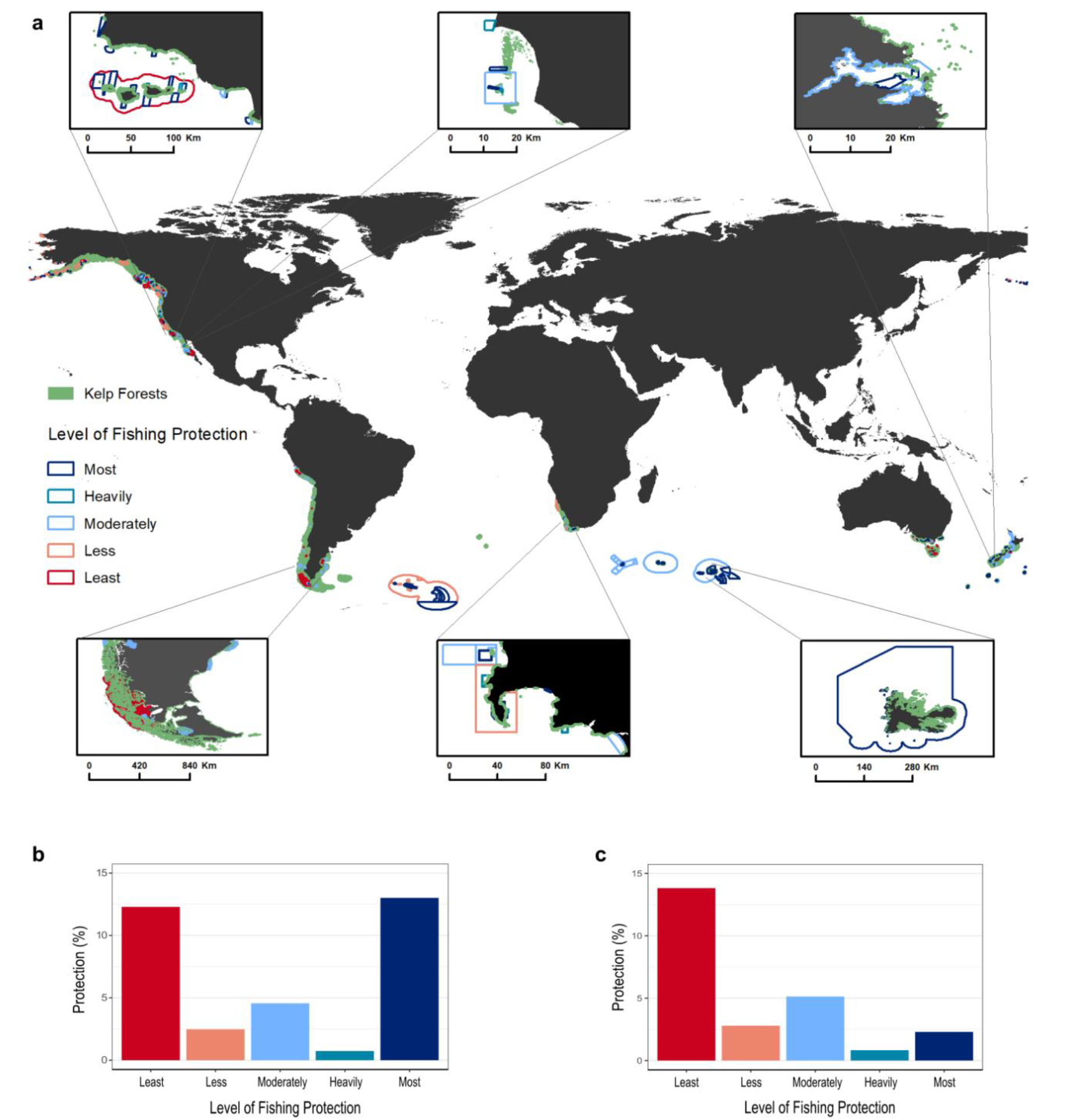
Global distribution of floating kelp forests and marine protected areas by categories of protection. **a** Global map of kelp forests and marine protected areas, we provide six fine-scale views. Starting from the top-left and moving clockwise: USA, Mexico, New Zealand, France, South Africa, and Chile and Argentina. Global protection (%) of kelp by category of protection **b** including all realms and **c** excluding the Southern Ocean realm. Protection categories are based on the Level of Fishing Protection (LFP)^38^ score assigned to each marine protected area. The scores are divided in three categories: Lightly protected (LFP score of “Least” and “Less”), moderately protected, and highly protected (LFP score of “Heavily” and “Most”).

The Arctic and the Temperate North Pacific are projected to be the most exposed to future MHWs under all climate scenarios, while Temperate South America will be the least exposed (Fig. 1b), matching the general spatial patterns in contemporary exposure. Overall, the pattern is very similar across SSP scenarios, with the northern hemisphere experiencing nearly twice the exposure to future MHWs than the southern hemisphere (Fig. 2b). However, some differences emerge. We found a difference in the latitudinal pattern of exposure between the northern and southern hemisphere. While in the northern hemisphere projections suggest a latitudinal pattern of increasing exposure to future MHWs from lower to higher latitudes, in the southern hemisphere, this pattern is reversed (Fig. 2b). For example, in the mid and long term and under all future scenarios for the northern hemisphere, the Eastern Bering Sea and the Gulf of Alaska are projected to become the most exposed ecoregions, while the southern California Bight becomes the least exposed (Fig. 2a and Supplementary Fig. 3), albeit with elevated levels of MHW exposure relative to the present. In contrast, in the southern hemisphere lower latitude ecoregions such as Cape Howe and Humboldtian are projected to be the most exposed to future MHWs while remote islands in high latitudes and ecoregions such as the Channels and Fjords of Southern Chile will be the least exposed.

### Global protection status of kelp forests

Globally, more than 33.1% of kelp forests are protected by MPAs, of which 13.7% are highly protected (the most effective type of MPAs), 4.6% are moderately protected, and 14.8% are in less-protected MPAs (Fig. 3a,b and 4a). However, most of the effective protection for kelp forests is in remote islands in the Southern Ocean realm, and when excluding these areas, only 2.8% of the global kelp forests are highly protected from fishing activities (Fig. 3c). At the country level, France has placed all their kelp forests within highly protected MPAs (Fig. 4a,b) and is the only country that meets the current 30% effective representation target5^7^. New Zealand, South Africa, Canada, Australia, and the USA have at least 10% of their kelp forests highly protected (Fig. 4a,b). However, this protection is in their overseas territories in remote islands for all of France (Kerguelen and Crozet Islands) and much of New Zealand, South Africa, and Australia. Australia has only 2.7%, New Zealand 3.2%, and South Africa 9.5% of their continental kelp forests highly protected. Mexico and Great Britain have provided effective protection for less than 2% of their kelp forests, Chile less than 0.02%, and Peru, Argentina, and Namibia none.

**Figure 4.**
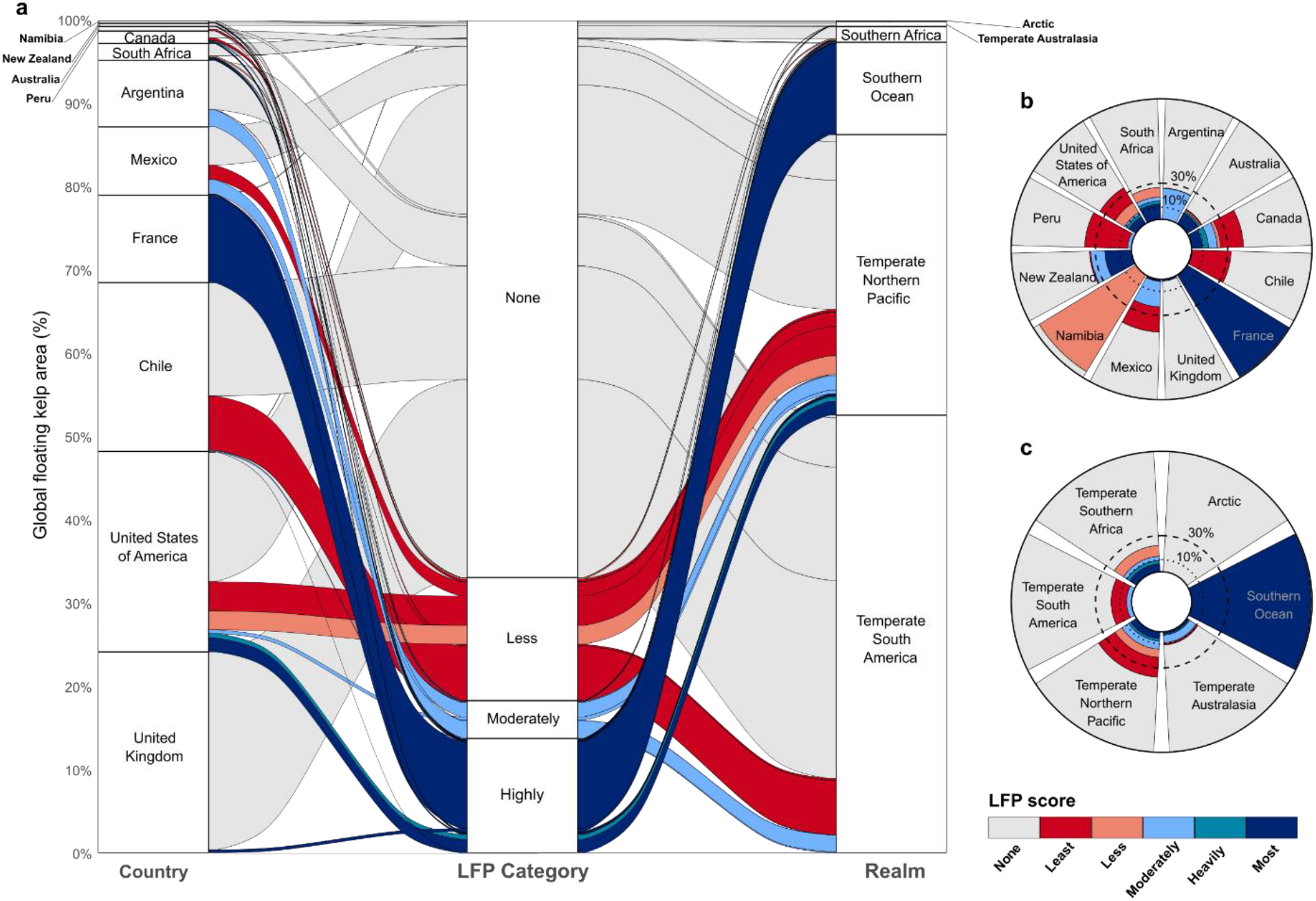
Global status and distribution of floating kelp forest protection. **a** Alluvial diagram with the distribution and protection of kelp by country and realm (% of total area), and radial plots showing percentage protection of kelp at the level of **b** country and **c** biogeographic realm. The dotted and dashed lines show the old 10%^2^ and the current 30%^7^ effective protection targets. Note that we included the Malvinas/Falkland Islands as part of the United Kingdom territory, although we acknowledge that Argentina has ongoing legal claims for their sovereignty.

Of the world’s biogeographic realms, the Southern Ocean has 99.9% of its kelp forests within highly protected MPAs, while all other realms have less than 10%. However, at least 10% of kelp forests are protected in some form of MPA in all realms, except for the Arctic, where the area of surface-canopy forming kelp is minimal and no kelp forests are protected under any category (Fig. 4c). At the ecoregional level, only 9 ecoregions have met the old 10% effective representation targets^2^ for kelp forests within highly protected MPAs, all in remote islands except for the Northern California ecoregion (Supplementary Fig. 4). Overall, 47.2% of ecoregions have less than 10% of the kelp forests protected, regardless of the MPA type. Only one nation, one realm, and 25% of ecoregions (all remote islands) meet the new 30% target for effective representation^7^ for kelp forests.

### Ecoregional future marine heatwave threats and protection status

Kelp forests within the ecoregions that are most threatened by projected MHWs and currently have low levels of effective protection (highly protected) include the Bering Sea (none protected), the Gulf of Alaska (0.6%), the North American Fjordlands (2.5%), the Puget Through (0.09%), and the Oregon to Vancouver ecoregions (2.4%) (Figure 5a,b and Supplementary Fig. 5 and 6). Northern California is the only ecoregion projected to be highly threated by MHWs where at least 10% of kelp forests are inside highly protected MPAs. In contrast, eight ecoregions that have all their kelp forests inside highly protected MPAs will face low to intermediate threats from projected MHWs under the SSP2.4-5 scenario. These ecoregions are all located in remote islands of the Southern Ocean realm. When combining highly and moderately protected MPAs, the Patagonian Shelf and North Patagonian Gulfs ecoregions have at least 30% of their kelp forests protected and low exposure to MHWs (Fig. 5b).

**Figure 5.**
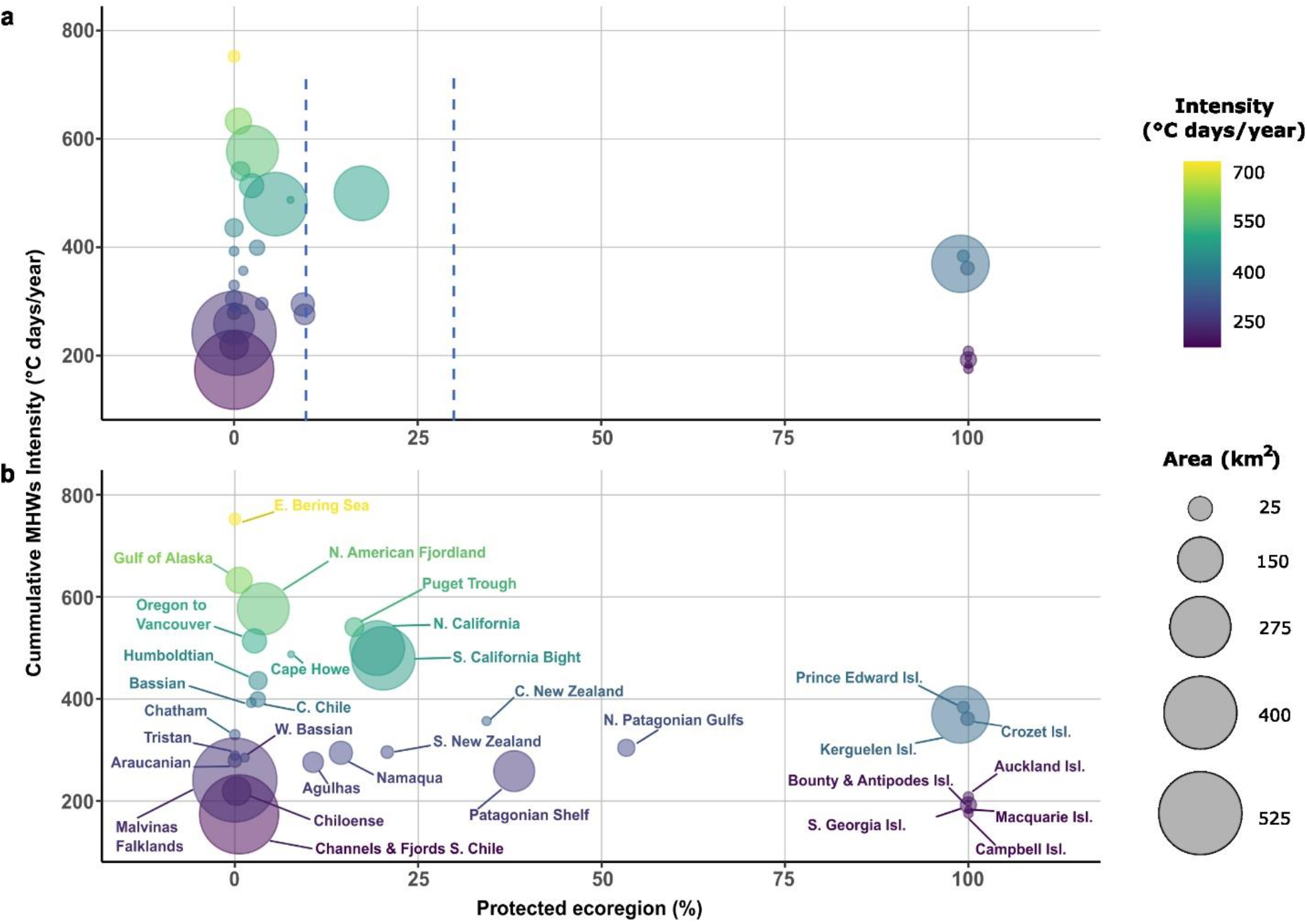
Relationship between threat posed by future marine heatwaves and level of protection for floating kelp forests. Scatterplots of mean future cumulative annual marine heatwave intensities for the midterm (2041-2060) under SSP2-4.5 and the amount of kelp forest **a** highly protected, **b** highly and moderately protected combined. The size of the bubble indicates the amount of kelp in each ecoregion. The dashed blue vertical lines represent the old 10%^**2**^ and the current 30%^**7**^ targets for effective protection.

## Discussion

We present the first global map of the protection status of floating kelp forests, which allowed us to identify escalating climate change threats and important conservation gaps for kelp forest ecosystems globally. Although one country and a few ecoregions are meeting current international protection targets^7^ for kelp forests, many of these MPAs are in remote islands with low levels of exposure to contemporary and projected MHWs and few non-climatic threats^39^. When kelp forests in remote islands are excluded, less than 3% of kelp forests are inside highly restrictive MPAs — no-take marine reserves— the most effective type of MPA for conserving biodiversity^1,40^ and for enhancing climate resilience^26-28,30,41-43^. Thus, current global protection does not adequately account for anthropogenic pressures on kelp forest ecosystems. It is concerning that the kelp forests most exposed to current and projected MHWs have minimal protection, which suggests that their resilience is likely being compromised. Therefore, to achieve international conservation commitments and climate adaptation goals, most countries and ecoregions require additional investments to increase the area of kelp forests that are effectively protected. This presents a unique opportunity for designing and implementing climate-smart MPAs^24^.

Our study reveals that marine heatwaves will increasingly threaten kelp forests under all projected SSP scenarios and time frames. If greenhouse emissions are not mitigated, kelp forests could be exposed to >16 times the magnitude of contemporary exposure under extreme scenarios by the end of the century. That represents an increase of 2–5°C in average ocean temperatures, which in some regions may permanently surpass physiological tolerances of kelp forests, impact their distribution, restructure associated ecological communities and impact the livelihood of local human communities^4,14,16,18,19,44-47^. Kelp forests near their current warm distribution limit will likely be the most affected and subject to range contractions^14,45,48,49^. Predicting whether MPAs can provide resilience to kelp forest ecosystems under such extreme and persistent changes is challenging. However, for less-extreme emission scenarios that track current mitigation policies^50,51^, the magnitude of exposure to future MHWs will be two times lower than for extreme scenarios. Under these conditions, it is more likely that marine reserves can support the resilience of kelp forests and enhance their adaptive capacity.

MPAs cannot directly mitigate the impacts of MHWs that surpass the physiological thresholds of kelp forests; however, they can minimize other non-climatic threats, such as overfishing and habitat destruction, thereby promoting the recovery of kelp forests following MHWs. For example, after the 2014–2016 MHWs in the northeast Pacific Ocean, urchins overgrazed kelp forests and caused many of them to collapse into less biodiverse ecosystems^15,22^. However, highly protected MPAs have prevented kelp forest collapse and have provided resilience to climate impacts by facilitating recovery of overfished predators that control urchin populations^29,30^. MPAs will likely not be enough to support the persistence of kelp forests, given the magnitude of future climate threats reported here, so other climate-adaptation strategies will be necessary, particularly for areas of high exposure to future MHWs, such as the Bering Sea, the Gulf of Alaska and the North American Fjordlands. These strategies include identifying and protecting climate refugia, restoring degraded kelp, identifying genetically resilient kelp stocks, and managing other anthropogenic impacts not mitigated by MPAs^15,52^.

We identified areas that will likely act as climate refugia —projected to be less exposed to future MHWs— where kelp forests are likely to persist^9,10,24,53^. We found that although many ecoregions with potential climate refugia have all their kelp forests protected inside MPAs, the Southern Fjordlands of Chile and the Malvinas/Falklands ecoregions have no protection and account for >40% of the global distribution of kelp forests. These ecoregions emerge as priority areas for global conservation of kelp forests, and efforts are needed to secure their effective protection and representation^39^ before other non-climatic threats intensify and erode their resilience.

It is important to note that our analysis includes only surface canopy kelp forests, thus excluding other kelp species. There are > 120 laminarian kelp species, of which only three of the largest kelp species form extensive floating canopies that can be detected by remote sensing. Our estimates likely represent overall kelp distribution and protection in regions where floating kelps co-exist with other kelp species. However, some areas included in our study and other nations and regions not included here have extensive subsurface kelp forests. Given the limitations in detecting subsurface canopy kelp species, they are likely less-well represented here than floating kelp species. This is a substantial gap for kelp conservation and an avenue for novel technologies and research^54^ to address associated needs, as these subsurface kelps also support diverse and productive ecosystems^13,55^ and human livelihoods^20^. We also note that our compiled map may overestimate or underestimate floating kelp coverage for those regions where validation of globally robust classifier maps is presently incomplete (e.g., Canada, Chile, New Zealand) and has been supplemented using a global map^35^. Therefore, the coverage of floating kelp reported here need to be carefully used for those regions and updated as new information becomes available.

Our analysis uses the distribution of present surface canopy kelp, and it does not account for range contractions or expansions of kelp forests that are projected under climate scenarios^45,46^. Integrating future range shifts of kelp species and associated biodiversity under climate scenarios could guide the identification of climate-smart priority areas for kelp forest conservation^24^. Finally, the MPA dataset used here has some limitations regarding the quantification of protection. For example, it does not account for other human activities that MPAs can manage (e.g., mining, dredging) or indicators of management efficiency (e.g., budget capacity, stage of establishment)^1^ that need to be included to ensure MPAs are effectively protecting ecosystems^56^. Therefore, including such information will likely decrease the coverage of kelp forests within highly protected MPAs. However, a comprehensive dataset of protection effectiveness is currently unavailable for all countries and MPAs (e.g., https://mpatlas.org/), and to date, Protected Seas^38^ is the most complete database available to assess the level of restriction inside MPAs.

## Conclusions

Kelp forests remain largely excluded from most international conservation policies^8,57^, despite their enormous contribution to earth’s biodiversity^12,13^ and provisioning of ecosystem services^20^. Nations have an opportunity to harness, protect, and restore kelp forests^58^, not only for their function as biogenic habitats and biodiversity hot spots^13^, but also to support their role in carbon sequestration and mitigation of climate change^59^. In addition, kelp forests provide food and support the livelihoods of millions of people worldwide^13,20^. As part of efforts to protect 30% of the oceans by 2030^7^, nations have an opportunity to explicitly include the representation of kelp forests in their national conservation policies. Where nations share ecoregions, transboundary management and coordination may also be needed^24^. However, given the immediate and escalating threats posed by climate change^14,16,44^ and other anthropogenic stressors, representation, though essential, may not be enough to secure the persistence of kelp forests. It is paramount that kelp forests are protected in each ecoregion, through representative, adequate, and well-connected networks of climate-smart MPAs that consider additional climate adaptation strategies^24^.

## Methods

### Mapping kelp forests

We compiled existing regional and national datasets of surface-canopy forming kelp derived using remote sensing observations (Supplementary Table 1). We compiled regional estimates of kelp canopy derived from up to four Landsat sensors: Landsat 5 Thematic Mapper (1984–2011), Landsat 7 Enhanced Thematic Mapper+ (1999–present), Landsat 8 Operational Land Imager (2013–present), and Landsat 9 Operational Land Imager-2 (2021-present). The applicable Landsat observations have pixel resolutions of 30 × 30 m and repeat times of 16 days (8 days since 1999 in most years because two Landsat sensors were operational). Classification of floating kelp canopy was derived by applying a globally robust random forest classifier to individual Landsat scenes^60^. The compiled datasets include minor differences in methodologies and time periods, but they all cover approximately over 30 years (1984 onwards) (Supplementary Table 1). Kelp maps were created by compositing observations of kelp presence across this time series. The maps include most of the USA (California, Oregon, parts of Washington, and parts of Alaska) and all of Mexico, Peru, and Argentina (available at https://kelpwatch.org/)^60^, most of the United Kingdom^61^ (Malvinas/Falkland Islands), and most of Australia^23^ (Tasmania). We included the Malvinas/Falkland Islands as part of the United Kingdom territory, although we acknowledge that Argentina has ongoing legal claims for their sovereignty.

We then used existing maps for South Africa^62^ and an empirical global map^35^ derived, both, from Sentintel-2 satellite data for the areas where the Landsat maps are not available. For the global empirical map, kelp area was calculated through a band-difference threshold algorithm validated using ground observations of *Macrocystis pyrifera* forests with high confidence in South America^33^. This method averages all the available images from the Sentinel-2 satellite sensor from 26 June 2015 to 23 June 2019 to create a cloud-free mosaic. It then applies band-difference thresholds to identify pixels likely containing floating kelp canopy and a land mask using global digital elevation models (ALOS and SRTM), discarding topography with elevation >0 m. The global map has not been validated for all areas and has some detection caveats; thus, region-specific uncertainties are unknown. For this reason, we excluded pixels that fell within a 30-m buffer relative to the coastline because the global map does not distinguish between intertidal green algae and floating kelp forests. We also masked those pixels that overlie estuaries, as this can also be a source of false positives. See Supplementary Table 1 for sources of information used to apply filters and masks. All kelp datasets were converted from coordinates to shapefiles with ArcMap10.8 using the World Geodetic System 1984 (WGS84). The final kelp habitat map includes any pixel that the satellite detected kelp at any point in the time series.

### Exposure of kelp forests to contemporary and future marine heatwaves

We estimated the expected threat of climate change to kelp forests by calculating historical and projected cumulative annual MHW intensities. Marine heatwaves are periods during which temperature exceeds the 90^th^ percentile of temperatures seasonally during a baseline period and last for at least five consecutive days^63^. To quantify the magnitude of present-day MHWs, we used the NOAA 0.25°-resolution Optimum Interpolation Sea-Surface Temperatures (OISST)^64^ dataset (1982-present).

We also considered MHW characteristics using SST outputs from each of 11 Coupled Model Intercomparison Project Phase 6 (CMIP6; Supplementary Table 6) Earth System models (ESMs) re-gridded to 0.25° resolution using bilinear interpolation in CDO (Climate Data Operators). We selected three climate scenarios generated under the IPCC Shared Socio-Economic Pathways (SPPs)^36^: SSP1-2.6, SSP2-4.5, SSP5-8.5. SSP1-2.6 represents an optimistic scenario with a peak in radiative forcing at ∼3 W m^-2^ by 2100 (approximating a future with 2°C of warming relative to the pre-industrial temperatures). SSP2-4.5 represents an intermediate mitigation scenario with radiative forcing stabilized at ∼4.5 W m^-2^ by 2100 (approximating implementation of current climate policies, resulting in 2.7°C of warming by 2100). SSP5-8.5 represents an extreme counterfactual climate scenario with a continued increase in greenhouse gas emissions with radiative forcing reaching 8.5 W m^-2^ by 2100 and rising after that. We bias-corrected the SST dataset from each ESM using the delta method (see^65^). This method ensures that the mean SST for each ESM was the same as that for the corresponding NOAA 0.25°-resolution OISST data for the reference period 1983–2014. We then determined which grid cells overlayed with kelp forests, and when the kelp cell had no corresponding SST data for the ESM models (because ESMs have relatively coarse resolution), we filled the cell using the inverse-distance-weighted mean of surrounding cells.

We then used the R package *heatwaveR*^*66*^ to estimate historical (1983–2020) and projected (2021– 2100) cumulative annual MHW intensity (ºC days) for each pixel using a baseline climatology of 1983–2012. Note that although we used OISST data to quantify contemporary MHW intensities, we used corresponding data from each ESM’s historical run for the period 1983–2014 when quantifying projected MHW intensities. This meant that we used ESM data in the baseline period instead of the OISST data, which ensured that inter-ESM skill in representing variability was faithfully captured. We used annual cumulative intensities because they are a good indicator of the exposure of kelp forests to warm anomalies^21,24^. We then estimated the median cumulative annual MHW intensity for each grid cell for the contemporary (2000–2021) period and across the 11 ESMs for the near-(2021–2040), mid- (2041–2060), and long-term (2081-2100) for each SSP and grid cell. Finally, we summarized trends in MHWs at the level of biogeographic realms and ecoregions^37^ by conducting a spatial overlay (following the same approach as in the next sections).

### Marine protected areas: level of fishing restriction

We obtained the spatial boundaries of MPAs using two different sources of information for the countries that have surface-canopy forming kelp forests. First, we downloaded MPA boundaries from official country-level agencies (Supplementary Table 1). We undertook extensive searches to ensure that we used the most updated official information, as global datasets can be less comprehensive at the country-level. We then categorized each MPA based on the level of restrictions to extractive activities. We used the Level of Fishing Protection (LFP) score obtained from Protected Seas^38^ (https://protectedseas.net/). This database scores MPAs based on fishing restrictions on a scale of 1–5 scale (1 = Least restricted, 2 = Less restricted, 3 = Moderately restricted, 4 = Heavily restricted, 5 = Most restricted). Protected Seas further divides the scores into categories: an LFP score of 1–2 is categorized as less protected, 3 as moderately protected, and 4–5 as highly protected areas, the most effective type of MPA. Finally, we reviewed both country-level and Protected Seas datasets and, when needed, consulted country-level experts to ensure that all MPAs were included. We did not include other types of spatial closures and area-based measures that are not MPAs. For a few MPAs (34 of 817) that had no LFP score, we reviewed existing information and assigned a new score based on the fishing restrictions reported at the country level. We did not include other regulatory activities that MPAs can manage (e.g., mining, dredging, anchoring) or indicators of management efficiency (e.g., enforcement capacity, budget capacity, implementing management plan)^1^ because such datasets are not comprehensively available for all countries.

### Global kelp distribution and protection

To estimate the amount of kelp within each level of protection, we performed a spatial intersection of MPA types (LFP classification; 817 spatial features) and the global kelp forest distribution (428,400 spatial features). Spatial intersection is a computationally expensive operation, so avoiding trivial calculations can significantly improve performance. We therefore developed and implemented a nested, parallelized, and hierarchical intersection algorithm. The approach is “nested” because spatial layers are split based on national jurisdiction before performing the spatial intersection. The approach is “parallelized” because the country-level intersection operations can be performed across parallel computer cores. Finally, the approach is “hierarchical” because, even within a country, not all kelp forests may lie within an MPA and not all MPAs may contain kelp. We first use a simple and less computationally expensive spatial join to identify kelp forests and MPAs that do not overlap with each other and exclude them from the expensive intersection calculation. Kelp forests excluded in this step are categorized as “not protected”. Finally, we perform the spatial intersection between the kelp forests and MPAs that overlap. We then repeated this approach at the biogeographic realms and ecoregions as outlined by^37^. For all operations, we used unprojected coordinates (EPSG code 4326) that uses WGS84 datum and a spherical geometry engine (s2)^67^ via the sf package^68^ in R. Parallelization was done using the furrr and future^69^ package in R. We validated geometries throughout the pipeline using ast_make_valid in sf; any invalid geometries were removed.

Knowing the location and amount of kelp protected, we proceeded to calculate the total extent of kelp by country, biogeographic realm, and ecoregion, and by MPA category and LFP score. We also determined how much kelp was outside any protection. All spatial analyses were performed in R version 4.3.1 (2023-06-16)^70^ using a x86_64-apple-darwin20 platform running macOS Ventura 13.4.1 and using the sf package v1.0^68,71^ with GEOS 3.11.0, GDAL 3.5.3, and PROJ 9.1.0.

### Ecoregional marine heatwave threats under SSP2.4-5 and kelp representation

Our final analysis assessed the relationship between the threats posed by projected future MHWs to kelp forests and the amount (% area) protected in each ecoregion. We conducted this analysis at the ecoregional scale because, ideally, networks of MPAs should be established to protect the underlying biophysical processes that maintain species distribution and composition^24^. Areas with low values of projected future MHW intensities are potential climate refugia for kelp forests. For simplicity, our measure of threat is focused only on the average cumulative MHW intensity under one SSP for each timeframe. We used SSP2.4-5 as an intermediate climate scenario that reflects less extreme outcomes and has been proposed to inform climate adaptation and policy^50,51^. Because the patterns of threat for each ecoregion are similar across time frames (i.e., magnitude is the largest difference across times), we focus in the main text on the mid-term and include results of the other times in the Supplementary information. We report results most conservatively for highly protected kelp, and then also for highly and moderately protected kelp combined. We did not include less protected MPAs in this analysis because this type of MPA provides minimal to no protection to marine ecosystems from extractive activities^1^.

## Data availability

The remote-sensing kelp forest dataset is available at https://portal.edirepository.org/nis/mapbrowse?packageid=knb-lter-sbc.74.13, https://kelpwatch.org/map, and https://biogeoscienceslaboxford.users.earthengine.app/view/kelpforests. The marine protected area database is available at https://protectedseas.net/ upon request. All other data needed to evaluate the conclusions in the paper are present in the paper or its Supplementary information. The codes used for this project will be made available upon publication.

## Acknowledgments

N.A.-D., K.C., and T.B. acknowledge support from the NASA Ocean Biology and Biogeochemistry program (80NSSC21K1429). N.A.-D. acknowledges support from the Estate Winifred Violet Scott (Australia) for a research grant. F.M. and N.A-D acknowledges support from the National Science Foundation (2108566). N.A-D, F.M, and K.C acknowledge funding from the Lenfest Ocean Program (ID Number: 00036969). We are very thankful to the Protected Seas team for sharing the marine protected area dataset.

## Author contributions

N.A-D. conceived the study with inputs from J.V-D., D.S., F.M., and K.C. A.M-S, T.B., H.H, L.D., C.B., and K.C. provided remote-sensing floating kelp forest datasets. N.A.-D, J.V-D., and D.S. conducted analyses. N.A.-D led reviewing nation-level marine protected area database with the support of A.M-S., L.D., K.S., C.P., D.P, and C.L. N.A-D. led the writing of the manuscript with the support of D.S, J.V-D., F.M., and K.C. All authors contributed to reviewing and editing of the manuscript.

## References

1 Grorud-Colvert, K. et al. The MPA Guide: A framework to achieve global goals for the ocean. Science 373, eabf0861 (2021).

2 Diversity, C. o. B. COP 10 Decision X/2: strategic plan for biodiversity 2011–2020. (2010).

3 Maxwell, S. L. et al. Area-based conservation in the twenty-first century. Nature 586, 217–227 (2020).

4 Pecl, G. T. et al. Biodiversity redistribution under climate change: Impacts on ecosystems and human well-being. Science 355, eaai9214 (2017).

5 Pörtner, H.-O. et al. Overcoming the coupled climate and biodiversity crises and their societal impacts. Science 380, eabl4881 (2023).

6 Parmesan, C., Morecroft, M. D. & Trisurat, Y. Climate change 2022: Impacts, adaptation and vulnerability, GIEC, (2022).

7 CBD. Nations Adopt Four Goals, 23 Targets for 2030 in Landmark UN Biodiversity Agreement. (2023).

8 Valckenaere, J., Techera, E., Filbee-Dexter, K. & Wernberg, T. Unseen and unheard: the invisibility of kelp forests in international environmental governance. Frontiers in Marine Science (2023).

9 Arafeh-Dalmau, N. et al. Southward decrease in the protection of persistent giant kelp forests in the northeast Pacific. Communications Earth & Environment 2, 119 (2021).

10 Arafeh-Dalmau, N., Olguín-Jacobson, C., Bell, T. W., Micheli, F. & Cavanaugh, K. C. Shortfalls in the protection of persistent bull kelp forests in the USA. Biological Conservation 283, 110133 (2023).

11 Jayathilake, D. R. & Costello, M. J. Version 2 of the world map of laminarian kelp benefits from more Arctic data and makes it the largest marine biome. Biological Conservation 257, 109099 (2021).

12 Schiel, D. R. & Foster, M. S. The biology and ecology of giant kelp forests. (Univ of California Press, 2015).

13 Wernberg, T., Krumhansl, K., Filbee-Dexter, K. & Pedersen, M. F. in World seas: An environmental evaluation 57–78 (Elsevier, 2019).

14 Arafeh-Dalmau, N. et al. Extreme marine heatwaves alter kelp forest community near its equatorward distribution limit. Frontiers in Marine Science 6, 499 (2019).

15 Arafeh-Dalmau, N. et al. Marine heat waves threaten kelp forests. Science 367, 635–635 (2020).

16 Smale, D. A. Impacts of ocean warming on kelp forest ecosystems. New Phytologist 225, 1447–1454 (2020).

17 Cavanaugh, K. C., Reed, D. C., Bell, T. W., Castorani, M. C. & Beas-Luna, R. Spatial variability in the resistance and resilience of giant kelp in southern and Baja California to a multiyear heatwave. Frontiers in Marine Science 6, 413 (2019).

18 Smale, D. A. et al. Marine heatwaves threaten global biodiversity and the provision of ecosystem services. Nature Climate Change 9, 306–312 (2019).

19 Smith, K. E. et al. Socioeconomic impacts of marine heatwaves: Global issues and opportunities. Science 374, eabj3593 (2021).

20 Eger, A. M. et al. The value of ecosystem services in global marine kelp forests. nature communications 14, 1894 (2023).

21 Oliver, E. C. et al. Projected marine heatwaves in the 21st century and the potential for ecological impact. Frontiers in Marine Science 6, 734 (2019).

22 McPherson, M. L. et al. Large-scale shift in the structure of a kelp forest ecosystem co-occurs with an epizootic and marine heatwave. Communications biology 4, 298 (2021).

23 Butler, C. L., Lucieer, V. L., Wotherspoon, S. J. & Johnson, C. R. Multi-decadal decline in cover of giant kelp Macrocystis pyrifera at the southern limit of its Australian range. Marine Ecology Progress Series 653, 1–18 (2020).

24 Arafeh-Dalmau, N. et al. Integrating climate adaptation and transboundary management: Guidelines for designing climate-smart marine protected areas. One Earth 6, 1523–1541 (2023).

25 Peleg, O., Blain, C. O. & Shears, N. T. Long-term marine protection enhances kelp forest ecosystem stability. Ecological Applications 33, e2895 (2023).

26 Ziegler, S. L. et al. Marine protected areas, marine heatwaves, and the resilience of nearshore fish communities. Scientific Reports 13, 1405 (2023).

27 Olguin-Jacobson, C. et al. Recovery mode: Marine protected areas enhance the resilience of kelp species from marine heatwaves. bioRxiv, 2024.2005. 2008.592820 (2024).

28 Ling, S., Johnson, C., Frusher, S. & Ridgway, K. Overfishing reduces resilience of kelp beds to climate-driven catastrophic phase shift. Proceedings of the National Academy of Sciences 106, 22341–22345 (2009).

29 Eisaguirre, J. H. et al. Trophic redundancy and predator size class structure drive differences in kelp forest ecosystem dynamics. Ecology 101, e02993 (2020).

30 Kumagai, J. A. et al. Marine protected areas promote resilience of kelp forests to marine heatwaves by preserving trophic cascades. bioRxiv, 2024.2004. 2010.588833 (2024).

31 Cavanaugh, K. C. et al. A Review of the Opportunities and Challenges for Using Remote Sensing for Management of Surface-Canopy Forming Kelps. Frontiers in Marine Science, 1536 (2021).

32 Keppel, G. et al. Refugia: identifying and understanding safe havens for biodiversity under climate change. Global Ecology and Biogeography 21, 393–404 (2012).

33 Woodson, C. B. et al. Harnessing marine microclimates for climate change adaptation and marine conservation. Conservation Letters 12, e12609 (2019).

34 Keppel, G. et al. The capacity of refugia for conservation planning under climate change. Frontiers in Ecology and the Environment 13, 106–112 (2015).

35 Mora-Soto, A. et al. A high-resolution global map of Giant kelp (Macrocystis pyrifera) forests and intertidal green algae (Ulvophyceae) with Sentinel-2 imagery. Remote Sensing 12, 694 (2020).

36 O’Neill, B. C. et al. The roads ahead: Narratives for shared socioeconomic pathways describing world futures in the 21st century. Global environmental change 42, 169–180 (2017).

37 Spalding, M. D. et al. Marine ecoregions of the world: a bioregionalization of coastal and shelf areas. BioScience 57, 573–583 (2007).

38 Driedger, A. et al. Guidance on marine protected area protection level assignments when faced with unknown regulatory information. Marine Policy 148, 105441 (2023).

39 Jones, K. R. et al. The location and protection status of Earth’s diminishing marine wilderness. Current Biology 28, 2506–2512. e2503 (2018).

40 Edgar, G. J. et al. Global conservation outcomes depend on marine protected areas with five key features. Nature 506, 216–220 (2014).

41 Micheli, F. et al. Evidence that marine reserves enhance resilience to climatic impacts. PLoS One 7, e40832 (2012).

42 Jacquemont, J., Blasiak, R., Le Cam, C., Le Gouellec, M. & Claudet, J. Ocean conservation boosts climate change mitigation and adaptation. One Earth 5, 1126–1138 (2022).

43 Benedetti-Cecchi, L. et al. Marine protected areas promote stability of reef fish communities under climate warming. Nature Communications 15, 1822 (2024).

44 Wernberg, T. et al. Climate-driven regime shift of a temperate marine ecosystem. Science 353, 169–172 (2016).

45 Assis, J., Fragkopoulou, E., Gouvêa, L., Araújo, M. B. & Serrão, E. A. Kelp forest diversity under projected end-of-century climate change. Diversity and Distributions, e13837 (2024).

46 Filbee-Dexter, K., Wernberg, T., Fredriksen, S., Norderhaug, K. M. & Pedersen, M. F. Arctic kelp forests: Diversity, resilience and future. Global and planetary change 172, 1–14 (2019).

47 Mora-Soto, A. et al. Kelp dynamics and environmental drivers in the southern Salish Sea, British Columbia, Canada. Frontiers in Marine Science (2024).

48 Smale, D. A. & Wernberg, T. Extreme climatic event drives range contraction of a habitat-forming species. Proceedings of the Royal Society B: Biological Sciences 280, 20122829 (2013).

49 Assis, J. et al. Major shifts at the range edge of marine forests: the combined effects of climate changes and limited dispersal. Scientific Reports 7, 44348 (2017).

50 Burgess, M. G., Becker, S. L., Langendorf, R. E., Fredston, A. & Brooks, C. M. Climate change scenarios in fisheries and aquatic conservation research. ICES Journal of Marine Science, fsad045 (2023).

51 Hausfather, Z. & Peters, G. P. Emissions–the ‘business as usual’story is misleading. Nature 577, 618–620 (2020).

52 Eger, A. M. et al. Global kelp forest restoration: past lessons, present status, and future directions. Biological Reviews 97, 1449–1475 (2022).

53 Mora-Soto, A. et al. A Song of Wind and Ice: Increased Frequency of Marine Cold-Spells in Southwestern Patagonia and Their Possible Effects on Giant Kelp Forests. Journal of Geophysical Research: Oceans 127, e2021JC017801 (2022).

54 Jayathilake, D. R. & Costello, M. J. Version 2 of the world map of laminarian kelp benefits from more Arctic data and makes it the largest marine biome. (2021).

55 Duarte, C. M. et al. Global estimates of the extent and production of macroalgal forests. Global Ecology and Biogeography 31, 1422–1439 (2022).

56 Pike, E. P. et al. Ocean protection quality is lagging behind quantity: Applying a scientific framework to assess real marine protected area progress against the 30 by 30 target. Conservation Letters, e13020.

57 Arafeh-Dalmau, N. et al. Introducing the Seaweed Specialist Group of the IUCN Species Survival Commission. Oryx 58, 147–148 (2024).

58 Eger, A. M. et al. The Kelp Forest Challenge: A collaborative global movement to protect and restore 4 million hectares of kelp forests. Journal of Applied Phycology, 1–14 (2023).

59 Pessarrodona, A. et al. Carbon sequestration and climate change mitigation using macroalgae: a state of knowledge review. Biological Reviews 98, 1945–1971 (2023).

60 Bell, T. W. et al. Kelpwatch: A new visualization and analysis tool to explore kelp canopy dynamics reveals variable response to and recovery from marine heatwaves. PLoS One 18, e0271477 (2023).

61 Houskeeper, H. F. et al. Automated satellite remote sensing of giant kelp at the Falkland Islands (Islas Malvinas). PLoS One 17, e0257933 (2022).

62 Dunga, V. L. Mapping and assessing ecosystem threat status of South African kelp forests, Faculty of Science, (2020).

63 Hobday, A. J. et al. A hierarchical approach to defining marine heatwaves. Progress in Oceanography 141, 227–238 (2016).

64 Huang, B. et al. Improvements of the daily optimum interpolation sea surface temperature (DOISST) version 2.1. Journal of Climate 34, 2923–2939 (2021).

65 Schoeman, D. S. et al. Demystifying global climate models for use in the life sciences. Trends in Ecology & Evolution (2023).

66 Schlegel, R. W. & Smit, A. J. heatwaveR: A central algorithm for the detection of heatwaves and cold-spells. Journal of Open Source Software 3, 821 (2018).

67 Dunnington D P. E., Rubak E s2: Spherical Geometry Operators Using the S2 Geometry Library. R package version 1.1.6. (https://github.com/r-spatial/s2, http://s2geometry.io/, https://r-spatial.github.io/s2/, 2024).

68 Pebesma, E. J. Simple features for R: standardized support for spatial vector data. R J. 10, 439 (2018).

69 Bengtsson, H. A unifying framework for parallel and distributed processing in R using futures. arXiv preprint arXiv:2008.00553 (2020).

70 R Core Team, R. R: A language and environment for statistical computing. (2013).

71 Pebesma, E. & Bivand, R. Spatial data science: With applications in R. (CRC Press, 2023).

