## Supplementary Information for "Intensifying marine heatwaves and limited protection threaten global kelp forests"

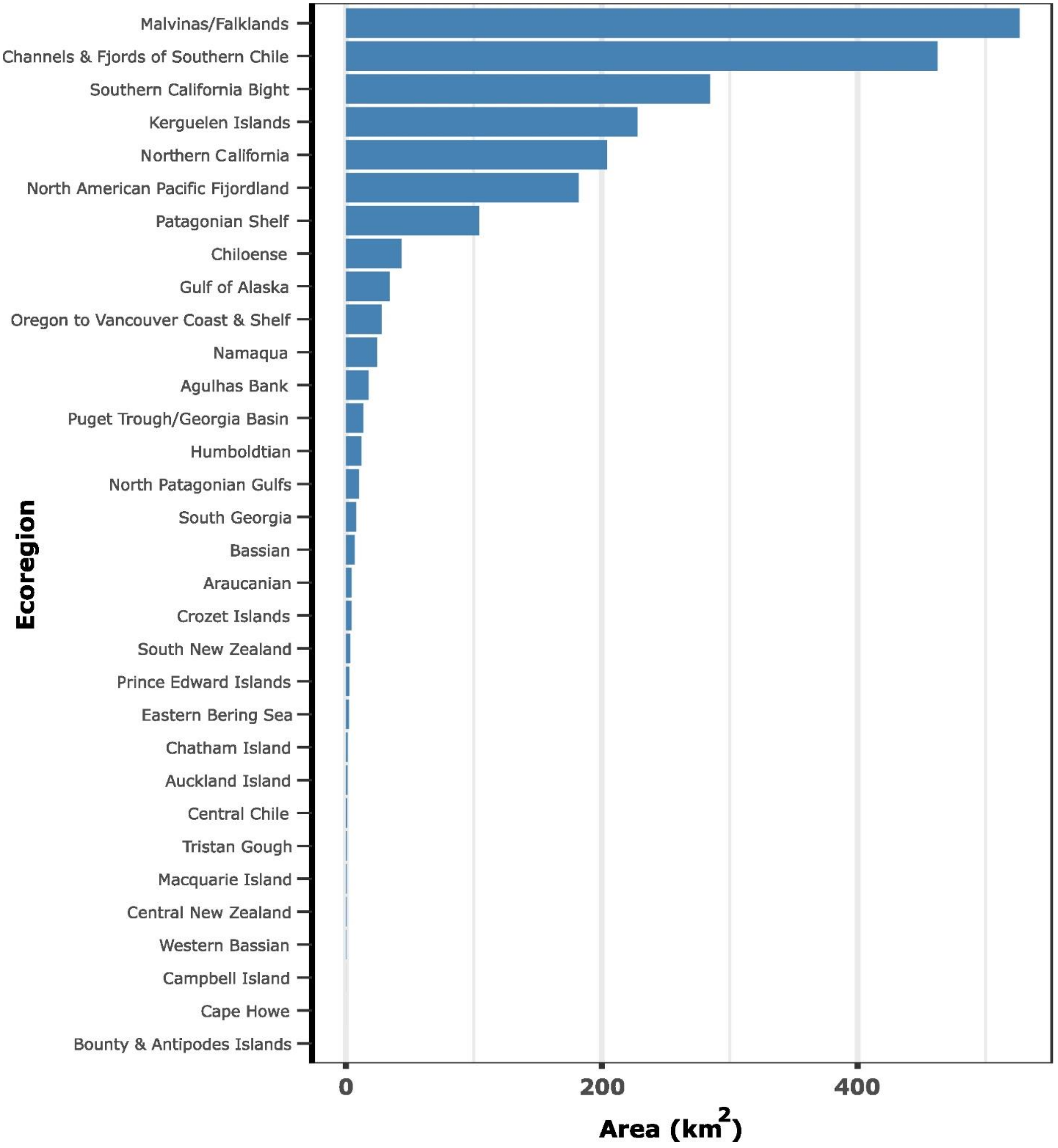


**Supplementary Figure 1.- Area of floating kelp forest (km$^2$) by ecoregion based on remote sensing maps.**

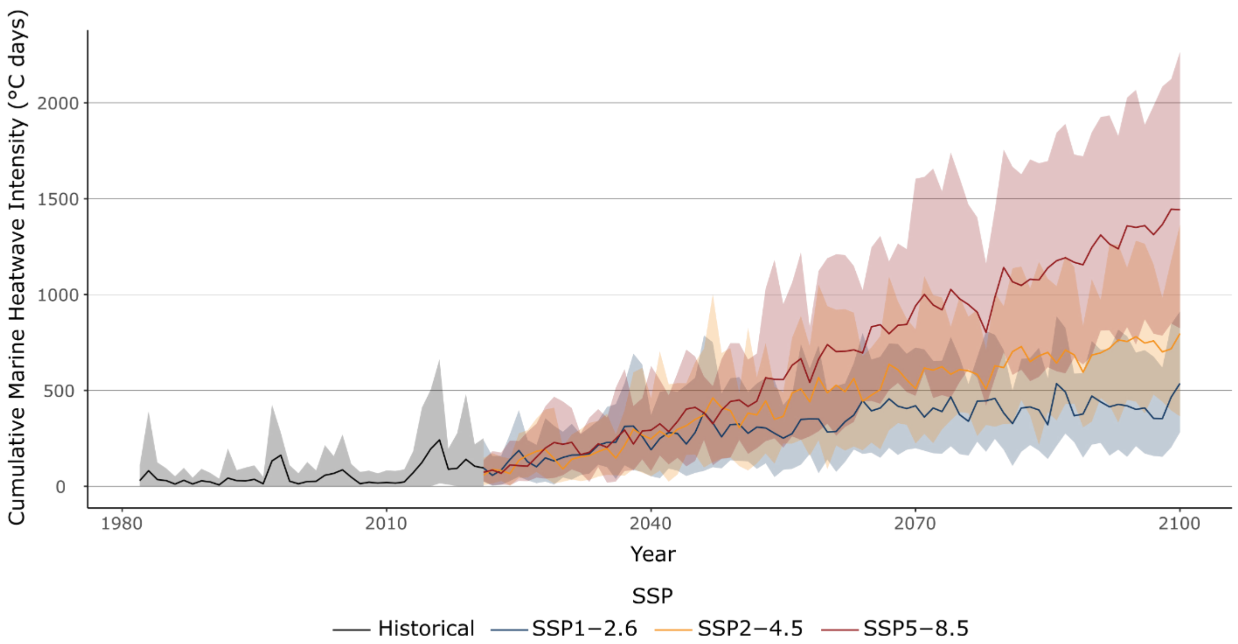


**Supplementary Figure 2.- Global exposure of floating kelp forests to contemporary and future marine heatwaves.** Global exposure of kelp forest to historical (1982-2020) and future cumulative annual MHW intensities (2021-2100) across three climate scenarios (SSP-1.26, SSP-2.45, SSP-5.85). The solid line shows the mean across ensemble medians for all pixels, and the shaded area represents the 5$^{th}$ and 95$^{th}$ percentiles.

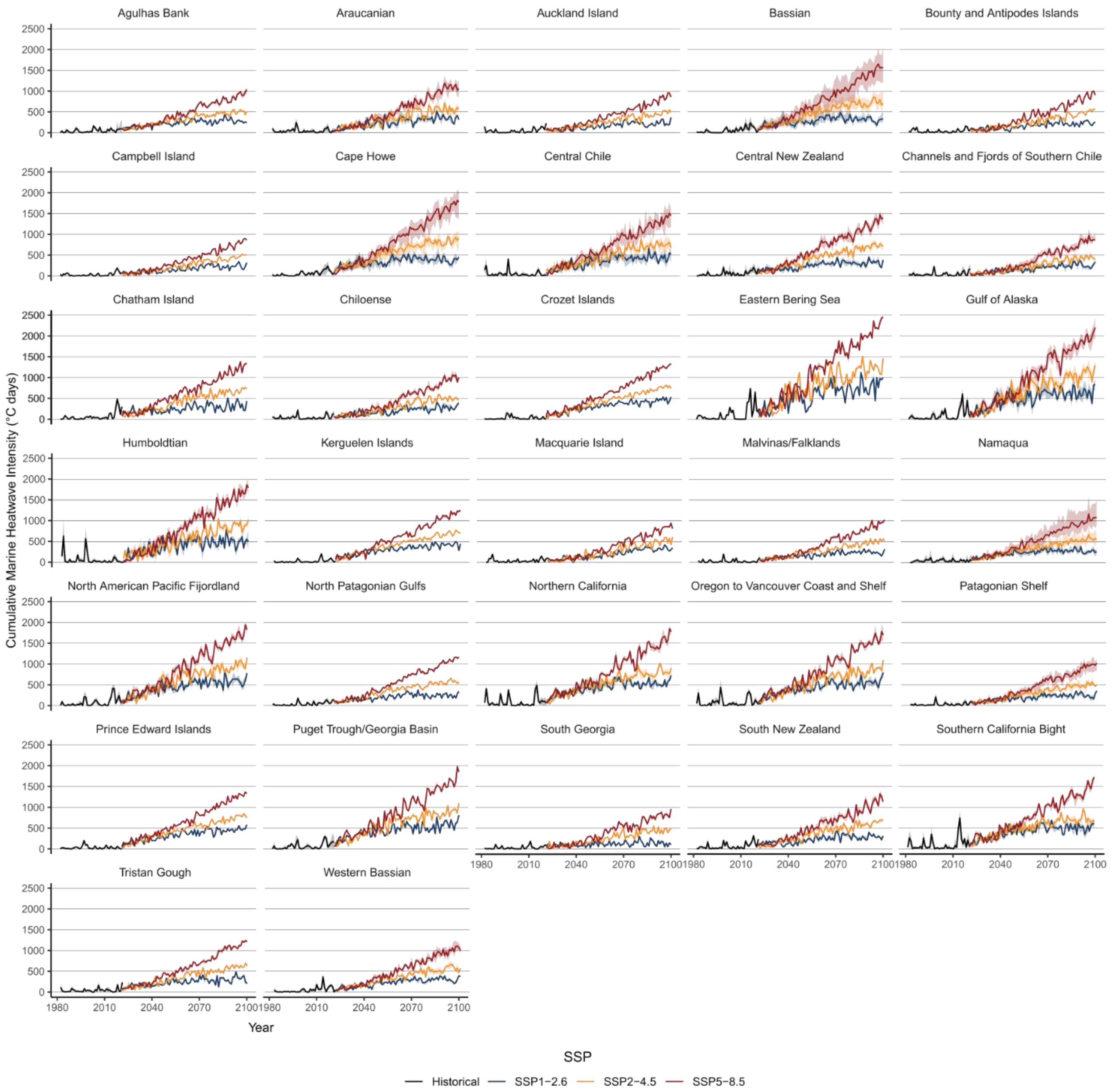


**Supplementary Figure 3.- Ecoregional exposure of floating kelp forests to contemporary and future marine heatwaves.** Ecoregion-specific exposure of kelp forest to historical (1982-2020) and future cumulative annual MHW intensities (2021-2100) across three climate scenarios (SSP-1.26, SSP-2.45, SSP-5.85). The solid line shows the mean across ensemble medians for all pixels, and the shaded area represents the 5$^{th}$ and 95$^{th}$ percentiles.

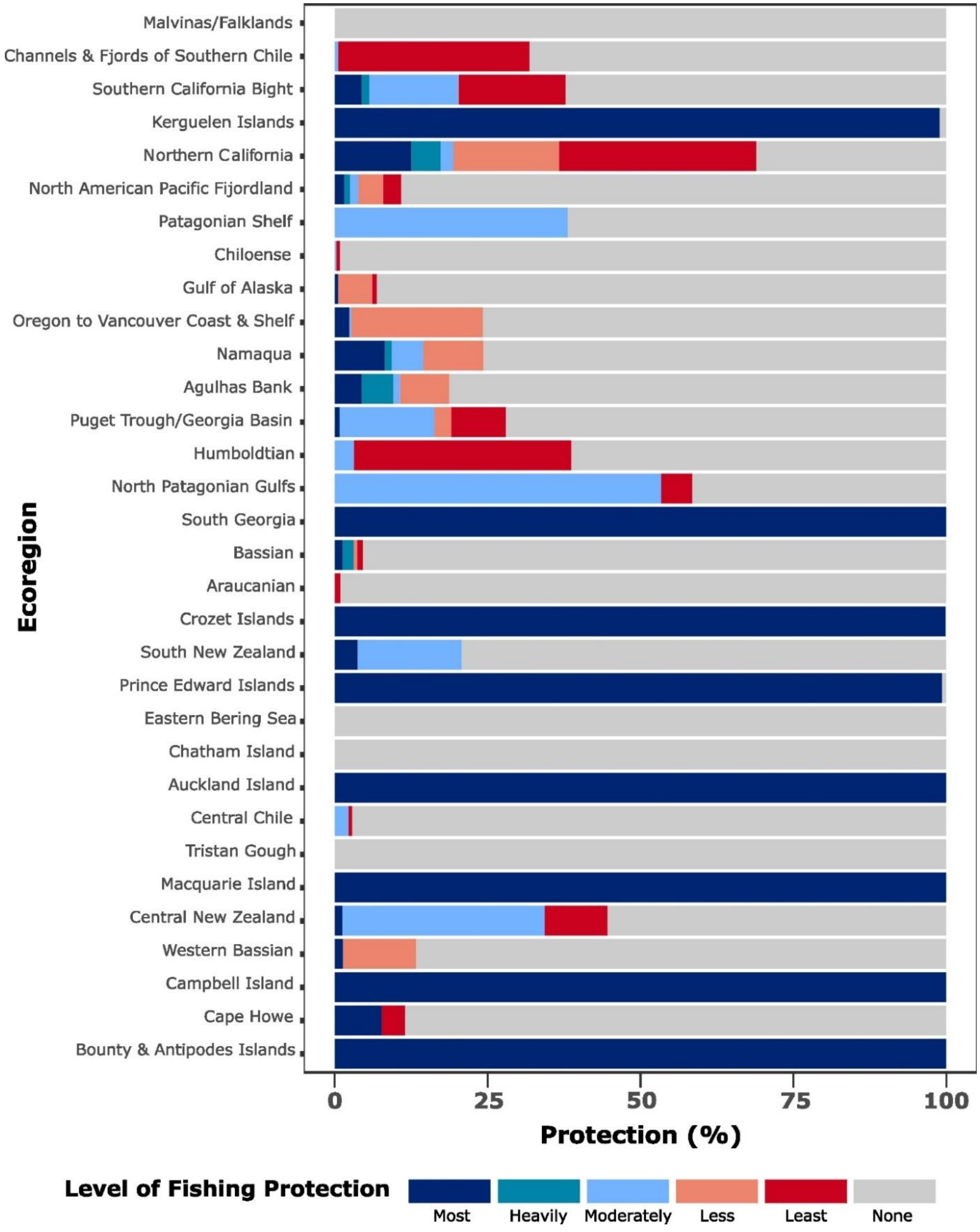


**Supplementary Figure 4.- Global protection of floating kelp forests for each ecoregion.** Percentage of kelp area protected in each ecoregion based on different levels of fishing restriction assigned to each marine protected area based on Protected Seas[1].

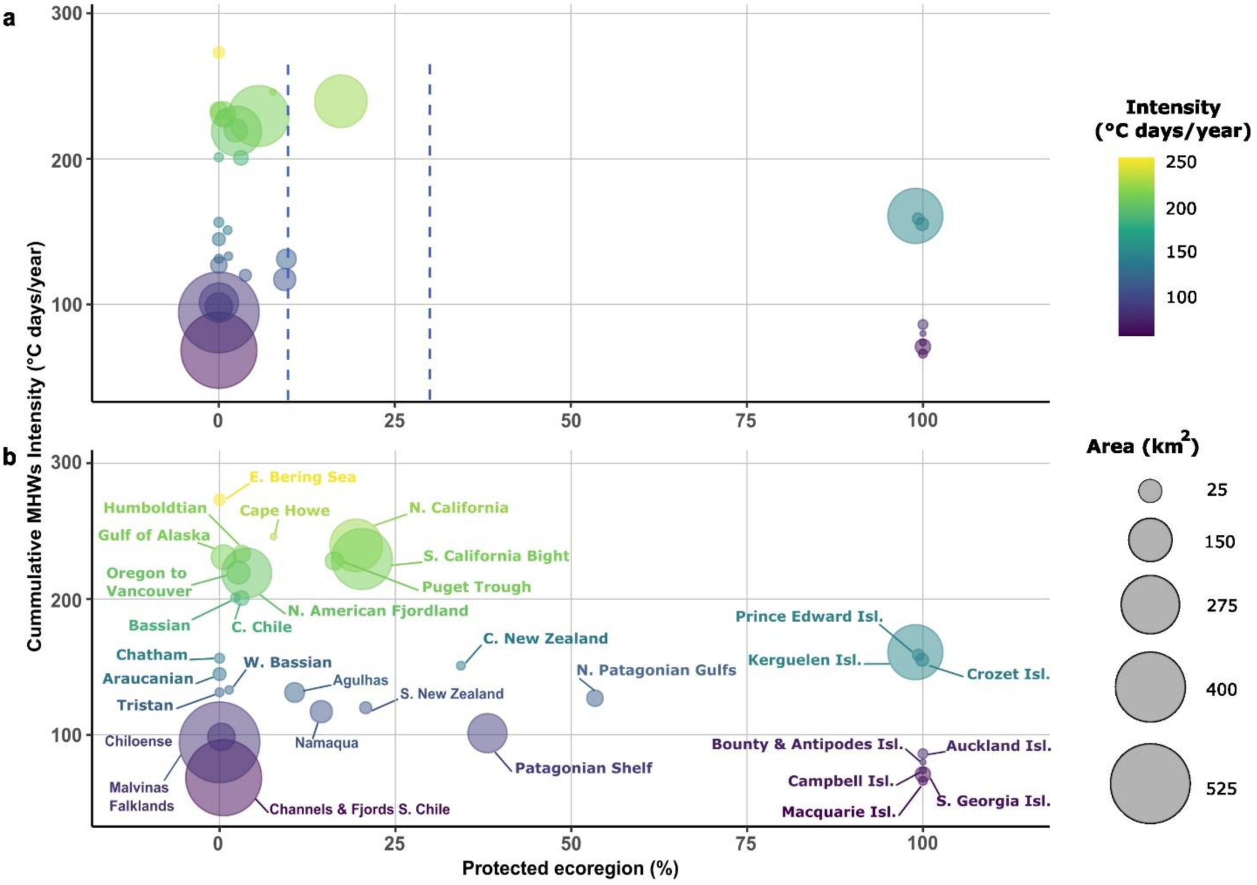


**Supplementary Figure 5.- Relationship between threat posed by future marine heatwaves and level of protection**. Scatterplots of mean future cumulative annual marine heatwave intensities for the near term (2021-2040) under SSP2-4.5 and the amount of kelp forest **a** highly protected, **b** highly and moderately protected combined. The size of the bubble indicates the amount of kelp in each ecoregion. The dashed blue vertical lines represent the old 10%[2] and the current 30%[3] targets for protection.

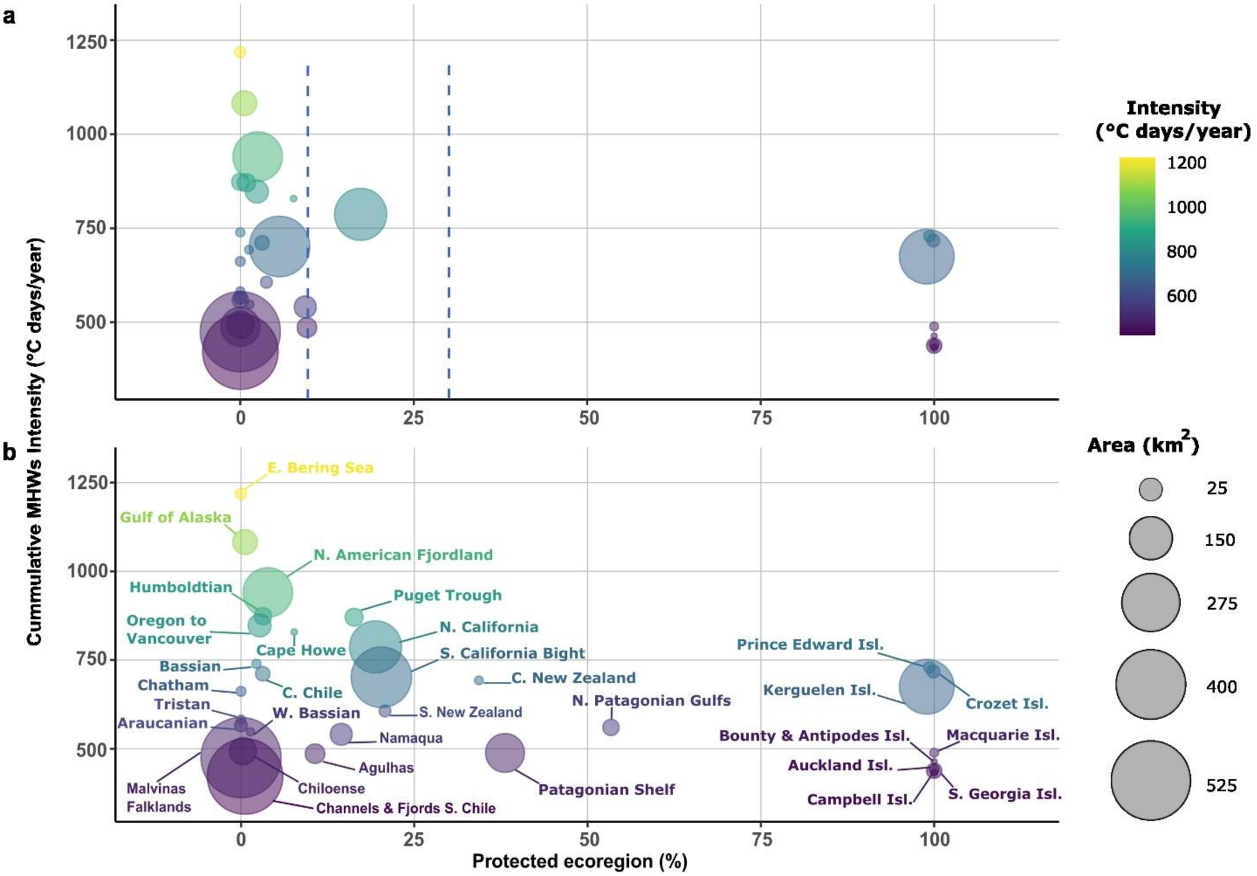


**Supplementary Figure 6.- Relationship between threat posed by future marine heatwaves and level of protection**. Scatterplots of mean future cumulative annual marine heatwave intensities for the long term (2081-2100) under SSP2-4.5 and the amount of kelp forest **a** highly protected, **b** highly and moderately protected combined. The size of the bubble indicates the amount of kelp in each ecoregion The dashed blue vertical lines represent the old 10%[2] and the current 30%[3] targets for protection.

**Supplementary Table 1.-** Data sources compiled to map the global distribution of floating kelp and marine protected areas (MPAs).

| County | Landsat images | Sentinel-2 images | Coastline | MPAs |
|---|---|---|---|---|
| USA | California, Oregon, western Washington (including the outer coast and most of the Strait of Juan de Fuca), and southern Alaska. Landsat images from 1984-2022[4,5] | For eastern and northern Washington, from Puget Sound, San Juan Islands, and the Strait of Georgia, and northern Alaska. Sentinel-2 images from 2015-2019[6] | Alaska<br>https://catalog.northslopescience.org/dataset/851/resource/5986cd56-eb50-44e0-915c-ec9ad71b0d0b<br><br>Washington:<br>https://shoreline.noaa.gov/data/datasheets/medres.html | Government (NOAA)<br>https://marineprotectedareas.noaa.gov/dataanalysis/mpainventory/ |
| Canada | | For British Columbia. Sentinel-2 images from 2015-2019. Includes all British Columbia[6]. | https://open.canada.ca/data/en/dataset/6c78fb2f-d23b-45b4-b3af-cc6f6cc4fff8 | Government (CPCAD)<br>https://www.canada.ca/en/environment-climate-change/services/national-wildlife-areas/protected-conserved-areas-database.html |
| Mexico | All kelp maps for the country are from Landsat images for 1984-2022[4] | | None | Government (CONANP)<br>http://sig.conanp.gob.mx/website/pagsig/info_shape.htm<br><br>Community-based MPAs and fishing refugees from COBI<br>https://cobi.org.mx/ |
| Peru | All kelp maps for the country are from Landsat images for 1984-present[4] | | None | Government (SERNANP)<br>https://geo.sernanp.gob.pe/visorsernanp/ |
| Chile | East Chilean Antarctic Province from Isla Nueva (Long ~ -66.4) to the Beagle Channel in the west (Long ~-69) from latitude -55.5 to the Chile-Argentina international border. Landsat images for 1984-2022[4] | Most kelp maps for the country are based on Sentinel-2 images from 2015-2019[6]. | https://www.marineregions.org/downloads.php | Government (SUBPESCA)<br>https://geoportal.subpesca.cl/portal/home/item.html?id=f1ece07b241740138fd07cb8404e328c |
| Argentina | All kelp maps for the country are from Landsat images for 1984-2022[4] | | None | Government (IGN)<br>https://www.ign.gob.ar/NuestrasActividades/InformacionGeoespacial/CapasSIG |

| | | | | |
|---|---|---|---|---|
| | | | | WCS Argentina https://ampargentina.org/ |
| Great Brittain | Falklands/Malvinas. Landsat images for 1984-2022[7] | For South Georgia and the South Sandwich Islands and Tristan da Cunha. Sentinel-2 images from 2015-2019[6] | Tristan da Cunha<br>https://www.marineregions.org/gazetteer.php?p=details&id=25511<br>South Georgia and the South Sandwich Islands<br>https://ramadda.data.bas.ac.uk/repository/entry/show?entryid=c1d83502-8799-4e3c-bdca-21db6a4405d4 | MPA Atlas https://mpatlas.org/mpaguide/ |
| Namibia | | All kelp maps for the country are from Sentinel-2 images (2015-2019)[6] | | MPA Atlas https://mpatlas.org/mpaguide/ |
| South Africa | | All kelp maps for continental South Africa are based on Landsat and Sentinel-2 images[8].<br>Prince Edwards Islands. Sentinel-2 images from 2015-2019[6] | Prince Edwards<br>https://marineregions.org/gazetteer.php?p=details&id=8384 | Government (SAPAD)<br>https://egis.environment.gov.za/data_egis/data_download/current |
| France | | All kelp maps for the country are from Sentinel-2 images (2015-2019)[6] | Crozet: https://www.marineregions.org/gazetteer.php?p=details&id=25547<br>Kerguelen: https://is.muni.cz/www/jaosobne/KergE.html | MPA Atlas https://mpatlas.org/mpaguide/ |
| Australia | Tasmania coastline. Landsat images from 1986-2015[9] | South Australia, Victoria, and Macquarie Island.<br>Sentinel-2 images from 2015-2019[6] | South Australia and Victoria<br>https://ecat.ga.gov.au/geonetwork/srv/eng/catalog.search#/metadata/61395<br>Macquarie Island https://data.gov.au/dataset/ds-dga-e4c01f74-de78-4233-80f4-444dfdc7380b/details | Government (CAPAD) https://www.environment.gov.au/fed/catalog/search/resource/details.page?uuid=%7BAF4EE98E-7F09-4172-B95E-067AB8FA10FC%7D<br>Government Tasmania https://listdata.thelist.tas.gov.au/opendata/index.html#LIST_Marine_Nature_Reserves |
| New Zealand | | All kelp maps for the country are from Sentinel-2 images (2015-2019)[6] | https://data.linz.govt.nz/data/?q=coastline | https://data.linz.govt.nz/layer/53564-protected-areas/ |

**Supplementary Table 2.-** Mean historical cumulative annual marine heatwave intensities (C° days) (± 95% confidence intervals) for the contemporary term (2021-2040), mid (2041-2060), and long term (2081-2100) based on OISST historical data.

| **Realm** | **Ecoregion** | **Contemporary** |
|---|---|---|
| Arctic | Eastern Bering Sea | 177.4 ± 15.1 |
| Temperate North Pacific | Gulf of Alaska | 136.7 ± 1.8 |
| | North American Pacific Fjordland | 106.8 ± 1.1 |
| | Northern California | 81.1 ± 2.2 |
| | Oregon to Vancouver Coast & Shelf | 78.3 ± 3.3 |
| | Puget Trough/Georgia Basin | 80.5 ± 6.2 |
| | Southern California Bight | 118.8 ± 3.5 |
| Temperate South America | Araucanian | 36.9 ± 1.9 |
| | Central Chile | 35.8 ± 3.76 |
| | Channels & Fjords of Southern Chile | 41.1 ± 1.2 |
| | Chiloense | 40.0 ± 1.3 |
| | Humboldtian | 30.1 ± 1.5 |
| | Falkland Islands (Malvinas) | 39.5 ± 0.9 |
| | North Patagonian Gulfs | 20.5 ± 0.7 |
| | Patagonian Shelf | 31.3 ± 2.7 |
| | Tristan Gough | 43.4 ± 2.2 |
| Southern Ocean | Auckland Island | 55.2 ± 3.7 |
| | Bounty & Antipodes Islands | 43.6 ± 4.3 |
| | Campbell Island | 44.3 ± 1.1 |
| | Crozet Islands | 25.28 ± 0.5 |
| | Kerguelen Islands | 39.5 ± 0.9 |
| | Macquarie Island | 43.5 ± 6.3 |
| | Prince Edward Islands | 23.7 ± 4.3 |
| | South Georgia | 35.7 ± 0.8 |
| Temperate Southern Africa | Agulhas Bank | 60.9 ± 5.9 |
| | Namaqua | 44.8 ± 3.8 |
| Temperate Australasia | Bassian | 90.5 ± 5.6 |
| | Cape Howe | 89.9 ± 18.6 |
| | Central New Zealand | 68.7 ± 4.3 |
| | Chatham Island | 93.9 ± 5.4 |
| | South New Zealand | 78.7 ± 1.2 |
| | Western Bassian | 85.1 ± 2.8 |

**Supplementary Table 3.-** Mean future cumulative annual marine heatwave intensities (C° days) (± 95% confidence intervals) for the near (2021-2040), mid (2041-2060), and long term (2081-2100) under SSP1-2.6.

| Realm | Ecoregion | Near-term SSP1.2-6 | Mid-term SSP1.2-6 | Long-term SSP1.2-6 |
|---|---|---|---|---|
| Arctic | Eastern Bering Sea | 299.6 ± 5.4 | 533.6 ± 6.9 | 786.7 ± 16.5 |
| Temperate North Pacific | Gulf of Alaska | 265.7 ± 1.5 | 476.0 ± 2.4 | 628.6 ± 4.7 |
| | North American Pacific Fjordland | 247.7 ± 1.0 | 440.3 ± 1.6 | 578.0 ± 1.5 |
| | Northern California | 253.4 ± 4.6 | 442.3 ± 3.8 | 537.5 ± 1.9 |
| | Oregon to Vancouver Coast & Shelf | 250.8 ± 5.1 | 418.2 ± 2.5 | 554.6 ± 4.0 |
| | Puget Trough/Georgia Basin | 258.8 ± 0.9 | 427.3 ± 0.6 | 569.6 ± 1.1 |
| | Southern California Bight | 224.9 ± 1.2 | 430.9 ± 3.7 | 507.4 ± 1.0 |
| Temperate South America | Araucanian | 120.9 ± 4.3 | 225.6 ± 8.2 | 346.4 ± 5.5 |
| | Central Chile | 178.4 ± 7.1 | 329.1 ± 17.3 | 447.5 ± 13.4 |
| | Channels & Fjords of Southern Chile | 75.3 ± 0.9 | 132.3 ± 0.9 | 258.2 ± 0.8 |
| | Chiloense | 98.6 ± 1.4 | 145.9 ± 2.6 | 273.9 ± 3.3 |
| | Humboldtian | 204.8 ± 6.5 | 425.1 ± 6.7 | 505.0 ± 8.6 |
| | Falkland Islands (Malvinas) | 97.7 ± 0.9 | 155.4 ± 0.8 | 245.3 ± 1.0 |
| | North Patagonian Gulfs | 117.3 ± 0.6 | 185.0 ± 1.1 | 243.2 ± 1.9 |
| | Patagonian Shelf | 107.9 ± 3.9 | 171.1 ± 2.6 | 250.5 ± 4.1 |
| | Tristan Gough | 142.6 ± 18.3 | 221.4 ± 14.7 | 310.6 ± 14.2 |
| Southern Ocean | Auckland Island | 88.4 ± 0.6 | 132.1 ± 3.0 | 219.3 ± 5.0 |
| | Bounty & Antipodes Islands | 93.5 ± 4.6 | 127.2 ± 5.8 | 220.0 ± 9.1 |
| | Campbell Island | 84.4 ± 1.1 | 125.5 ± 0.9 | 226.0 ± 2.1 |
| | Crozet Islands | 155.3 ± 1.9 | 284.5 ± 0.9 | 440.8 ± 1.5 |
| | Kerguelen Islands | 167.9 ± 1.4 | 288.3 ± 2.0 | 420.8 ± 2.7 |
| | Macquarie Island | 77.63 ± 3.4 | 141.9 ± 4.4 | 328.0 ± 8.9 |
| | Prince Edward Islands | 180.03 ± 1.1 | 305.1 ± 1.7 | 463.4 ± 2.8 |
| | South Georgia | 88.53 ± 0.9 | 97.3 ± 2.0 | 131.1 ± 2.3 |
| Temperate Southern Africa | Agulhas Bank | 131.3 ± 4.9 | 224.2 ± 3.4 | 287.0 ±1.4 |
| | Namaqua | 136.5 ± 5.3 | 230.1 ± 4.5 | 277.3 ± 3.5 |
| Temperate Australasia | Bassian | 211.9 ± 18.4 | 250.4 ± 16.5 | 312.7 ± 10.0 |
| | Cape Howe | 261.0 ± 46.7 | 327.4 ± 49.0 | 383.7 ± 41.5 |
| | Central New Zealand | 160.5 ± 5.0 | 222.4 ± 4.3 | 310.1 ± 3.6 |
| | Chatham Island | 177.9 ± 4.3 | 204.3 ± 2.0 | 311.6 ± 1.7 |
| | South New Zealand | 137.6 ± 2.7 | 188.1 ± 3.9 | 285.4 ± 1.5 |
| | Western Bassian | 124.2 ± 5 | 204.1 ± 3.7 | 310.3 ± 1.8 |

**Supplementary Table 4.-** Mean future cumulative annual marine heatwave intensities (C° days) (± 95% confidence intervals) for the near (2021-2040), mid (2041-2060), and long term (2081-2100) under SSP2-4.5.

| Realm | Ecoregion | Near-term SSP2.4-5 | Mid-term SSP2.4-5 | Long-term SSP2.4-5 |
|---|---|---|---|---|
| Arctic | Eastern Bering Sea | 273.1 ± 7.0 | 752.4 ± 20.5 | 1218.2 ± 17.1 |
| Temperate North Pacific | Gulf of Alaska | 230.8 ± 1.5 | 632.7 ± 4.1 | 1082.6 ± 6.8 |
| | North American Pacific Fjordland | 219.1 ± 1.0 | 576.8 ± 2.3 | 940.5 ± 3.6 |
| | Northern California | 239.6 ± 1.6 | 499.7 ± 2.3 | 787.0 ± 7.5 |
| | Oregon to Vancouver Coast & Shelf | 219.5 ± 2.6 | 513.7 ± 8.4 | 847.4 ± 6.2 |
| | Puget Trough/Georgia Basin | 228.1 ± 0.5 | 540.6 ± 0.5 | 870.8 ± 0.3 |
| | Southern California Bight | 229.4 ± 2.3 | 479.6 ± 6.0 | 701.5 ± 13.0 |
| Temperate South America | Araucanian | 144.7 ± 4.7 | 279.4 ± 7.7 | 565.8 ± 12.6 |
| | Central Chile | 201.1 ± 7.5 | 392.7 ± 19.0 | 739.0 ± 23.5 |
| | Channels & Fjords of Southern Chile | 68.5 ± 1.2 | 173.9 ± 2.0 | 422.0 ± 2.6 |
| | Chiloense | 98.5 ± 1.6 | 219.4 ± 3.0 | 493.2 ± 2.2 |
| | Humboldtian | 233.2 ± 7.1 | 436.0 ± 8.5 | 873.5 ± 3.3 |
| | Falkland Islands (Malvinas) | 94.5 ± 0.8 | 241.2 ± 1.0 | 475.0 ± 1.3 |
| | North Patagonian Gulfs | 127.0 ± 1.4 | 304.3 ± 2.9 | 559.9 ± 0.9 |
| | Patagonian Shelf | 101.3 ± 5.2 | 258.6 ± 8.5 | 488.1 ± 10.8 |
| | Tristan Gough | 131.4 ± 4.1 | 288.8 ± 12.0 | 582.3 ± 14.6 |
| Southern Ocean | Auckland Island | 86.2 ± 1.9 | 208.1 ± 2.8 | 441.2 ± 6.4 |
| | Bounty & Antipodes Islands | 80.0 ± 2.1 | 195.8 ± 8.8 | 464.0 ± 13.9 |
| | Campbell Island | 73.9 ± 0.1 | 182.6 ± 2.2 | 432.4 ± 2.9 |
| | Crozet Islands | 155.2 ± 2.3 | 361.3 ± 1.9 | 717.4 ± 2.2 |
| | Kerguelen Islands | 160.8 ± 1.2 | 369.4 ± 2.2 | 674.7 ± 2.7 |
| | Macquarie Island | 66.2 ± 6.2 | 175.5 ± 5.8 | 488.9 ± 11.2 |
| | Prince Edward Islands | 158.9 ± 0.4 | 383.9 ± 2.1 | 729.2 ± 3.9 |
| | South Georgia | 70.9 ± 0.9 | 192.4 ± 3.7 | 437.8 ± 4.2 |
| Temperate Southern Africa | Agulhas Bank | 131.2 ± 2.5 | 276.0 ± 6.5 | 486.0 ± 9.3 |
| | Namaqua | 117.1 ± 3.1 | 294.5 ± 9.9 | 540.6 ± 23.4 |
| Temperate Australasia | Bassian | 200.7 ± 16.2 | 399.2 ± 21.9 | 711.3 ± 25.9 |
| | Cape Howe | 245.9 ± 37.3 | 487.5 ± 64.9 | 828.9 ± 68.6 |
| | Central New Zealand | 151.0 ± 2.6 | 356.7 ± 5.7 | 692.6 ± 9.1 |
| | Chatham Island | 156.4 ± 3.0 | 329.9 ± 1.0 | 661.7 ± 5.3 |
| | South New Zealand | 119.9 ± 2.1 | 295.9 ± 4.2 | 606.5 ± 5.3 |
| | Western Bassian | 133.1 ± 4.3 | 284.9 ± 10.9 | 547.4 ± 20.6 |

**Supplementary Table 5.-** Mean future cumulative annual marine heatwave intensities (C° days) (± 95% confidence intervals) for the near (2021-2040), mid (2041-2060), and long term (2081-2100) under SSP5-8.5.

| Realm | Ecoregion | Near-term SSP5.8-5 | Mid-term SSP5.8-5 | Long-term SSP5.8-5 |
|---|---|---|---|---|
| Arctic | Eastern Bering Sea | 358.9 ± 26.5 | 870.4 ± 21.4 | 2042.8 ± 17.8 |
| Temperate North Pacific | Gulf of Alaska | 283.7 ± 2.5 | 718.8 ± 6.7 | 1830.9 ± 9.1 |
| | North American Pacific Fjordland | 280.8 ± 0.8 | 660.1 ± 2.1 | 1588.3 ± 3.9 |
| | Northern California | 276.9 ± 2.5 | 582.0 ± 1.6 | 1469.2 ± 11.4 |
| | Oregon to Vancouver Coast & Shelf | 278.6 ± 4.6 | 612.4 ± 9.6 | 1504.1 ± 10.5 |
| | Puget Trough/Georgia Basin | 291.0 ± 0.9 | 640.1 ± 1.3 | 1535.8 ± 1.3 |
| | Southern California Bight | 259.6 ± 2.8 | 574.7 ± 4.1 | 1352.8 ± 13.4 |
| Temperate South America | Araucanian | 122.4 ± 5.6 | 361.8 ± 13.7 | 1019.8 ± 14.7 |
| | Central Chile | 207.1 ± 11.9 | 538.5 ± 18.2 | 1269.4 ± 38.8 |
| | Channels & Fjords of Southern Chile | 71.8 ± 1.1 | 225.1 ± 2.4 | 764.9 ± 5.1 |
| | Chiloense | 94.6 ± 0.8 | 292.1 ± 3.1 | 886.0 ± 3.8 |
| | Humboldtian | 186.9 ± 8.6 | 624.5 ± 6.3 | 1551.1 ± 11.6 |
| | Falkland Islands (Malvinas) | 98.3 ± 1.3 | 276.1 ± 1.6 | 835.5 ± 4.1 |
| | North Patagonian Gulfs | 139.7 ± 0.6 | 404.1 ± 2.1 | 1014.8 ± 2.6 |
| | Patagonian Shelf | 105.8 ± 4.2 | 316.3 ± 11.3 | 874.0 ± 22.7 |
| | Tristan Gough | 149.7 ± 6.3 | 423.0 ± 15.6 | 1081.6 ± 6.0 |
| Southern Ocean | Auckland Island | 92.9 ± 1.8 | 284.9 ± 3.2 | 753.4 ± 5.7 |
| | Bounty & Antipodes Islands | 88.9 ± 3.9 | 279.3 ± 4.2 | 791.2 ± 15.8 |
| | Campbell Island | 89.7 ± 1.1 | 256.9 ± 4.4 | 722.4 ± 7.5 |
| | Crozet Islands | 176.1 ± 2.9 | 472.0 ± 1.0 | 1155.7 ± 2.4 |
| | Kerguelen Islands | 160.9 ± 1.6 | 468.1 ± 1.5 | 1089.4 ± 4.7 |
| | Macquarie Island | 77.1 ± 3.8 | 217.1 ± 11.3 | 742.7 ± 15.8 |
| | Prince Edward Islands | 203.7 ± 4.6 | 486.2 ± 1.7 | 1194.7 ± 7.8 |
| | South Georgia | 68.1 ± 1.9 | 253.1 ± 3.2 | 770.3 ± 6.4 |
| Temperate Southern Africa | Agulhas Bank | 140.3 ± 4.2 | 337.7 ± 11.3 | 838.2 ± 11.3 |
| | Namaqua | 152.9 ± 5.8 | 379.5 ± 18.3 | 959.6 ± 49.9 |
| Temperate Australasia | Bassian | 193.9 ± 16.2 | 551.1 ± 30.5 | 1405.4 ± 62.8 |
| | Cape Howe | 223.3 ± 37.0 | 623.2 ± 60.3 | 1583.4 ± 137.5 |
| | Central New Zealand | 174.2 ± 3.2 | 496.1 ± 7.7 | 1194.4 ± 15.2 |
| | Chatham Island | 165.0 ± 2.8 | 508.3 ± 3.1 | 1147.5 ± 6.8 |
| | South New Zealand | 144.6 ± 3.3 | 416.2 ± 6.5 | 1013.5 ± 12.1 |
| | Western Bassian | 109.9 ± 7.5 | 368.0 ± 15.6 | 956.9 ± 35.3 |

**Supplementary Table 6.-** Climate models with daily sea surface temperature used in this study.

| **Modelling Center** | **Institute ID** | **Model Name** |
|---|---|---|
| Commonwealth Scientific and Industrial Research Organisation, Australia | CSIRO | ACCESS-CM2<br>ACCESS-ESM1 |
| Beijing Climate Center, China Meteorological Administration | BC | BCC-CSM2-MR |
| Canadian Centre for Climate Modelling and Analysis | CCCMA | CanESM5 |
| Centre National de Recherches Meteorologiques/ Centre Europeen de Recherche et Formation Avancees en Calcul Scientifique | CNRM-CERFACS | CNRM-CM6-1<br>CNRM-ESM2-1 |
| EC-Earth consortium | EC-Earth | EC-Earth3 |
| Atmosphere and Ocean Research Institute (The University of Tokyo), National Institute for Environmental Studies, and Japan Agency for Marine -Earth Science and Technology | MIROC | MIROC6 |
| Meteorological Research Institute | MRI | MRI-ESM2-0 |
| Norwegian Climate Center | NCC | NorESM2-LM<br>NorESM2-MM |

### References


1. Driedger, A. *et al.* Guidance on marine protected area protection level assignments when faced with unknown regulatory information. *Marine Policy* **148**, 105441 (2023).
2. CBD. Strategic Plan for Biodiversity 2011–2020, Convention on Biological Diversity. Montreal., (2010).
3. Diversity, C. o. B. (Convention on Biological Diversity Montreal).
4. Bell, T. W. *et al.* Kelpwatch: A new visualization and analysis tool to explore kelp canopy dynamics reveals variable response to and recovery from marine heatwaves. *PLoS One* **18**, e0271477 (2023).
5. McPherson, M. L. *et al.* Large-scale shift in the structure of a kelp forest ecosystem co-occurs with an epizootic and marine heatwave. *Communications biology* **4**, 1-9 (2021).
6. Mora-Soto, A. *et al.* A high-resolution global map of Giant kelp (Macrocystis pyrifera) forests and intertidal green algae (Ulvophyceae) with Sentinel-2 imagery. *Remote Sensing* **12**, 694 (2020).
7. Houskeeper, H. F. *et al.* Automated satellite remote sensing of giant kelp at the Falkland Islands (Islas Malvinas). *PLoS One* **17**, e0257933 (2022).
8. Dunga, V. L. Mapping and assessing ecosystem threat status of South African kelp forests. (2020).
9. Butler, C. L., Lucieer, V. L., Wotherspoon, S. J. & Johnson, C. R. Multi-decadal decline in cover of giant kelp Macrocystis pyrifera at the southern limit of its Australian range. *Marine Ecology Progress Series* **653**, 1-18 (2020).
